# Mobility of Four Structural Regions Drives Isoform-Specific Properties of Plant LPOR

**DOI:** 10.1101/2024.10.21.619389

**Authors:** Michał Gabruk, Mateusz Łuszczyński, Katarzyna Szafran, Wiktoria Ogrodzińska, Brenda M. Rubenstein, Gabriel Monteiro da Silva

## Abstract

Light-dependent protochlorophyllide oxidoreductase (LPOR) is a photocatalytic enzyme in the chlorophyll (Chl) biosynthetic pathway that underwent duplications in angiosperms, resulting in the emergence of multiple isoforms across various plant species. The physiological roles of these LPOR homologs remained unclear, so we selected six plant species with different number of isoforms of the enzyme and characterized their properties in vitro.

Our findings revealed that these isoforms vary in their affinity for the reaction product, chlorophyllide (Chlide), as well as for NADPH under lipid-free conditions and in reaction mixtures supplemented with plant lipids. Additionally, we observed differences in their oligomerization behavior.

Our experimental approach generated a dataset comprising several hundred pairs of spectra, recorded before and after reaction-triggering illumination. This data was used to analyze the correlation between fluorescence emission maxima before and after photoconversion. The analysis showed that some isoforms rapidly release Chlide after the reaction, while others retain the pigment in the binding pocket, especially at high NADPH concentrations. These results suggest that LPOR isoforms differ in their rates of Chlide release and complex disassembly, potentially influencing the overall rate of the Chl biosynthetic pathway, even in mature leaves.

We further analyzed the flexibility of these isoforms using AlphaFold2 predictions, identifying four regions of the enzyme that are particularly mobile. Two of these regions are involved in pigment binding, while the other two play a role in oligomerization. Based on these findings, we propose a model of conformational changes that drive the formation of LPOR oligomers.

## Introduction

Light-dependent protochlorophyllide oxidoreductase (LPOR) is a photocatalytic enzyme in the chlorophyll (Chl) biosynthetic pathway^1,2^, crucial for forming the paracrystalline structure of prolamellar bodies (PLBs) in etiolated plants^3,4^. LPOR has been shown to remodel lipid membranes containing monogalactosyl diacylglycerol (MGDG), with the resulting structure depending on the dinucleotide bound alongside its substrate, protochlorophyllide (Pchlide), within the binding pocket. NADPH binding leads to the formation of ternary Pchlide:LPOR:NADPH complexes with photocatalytic properties^5^, which in darkness organize into filamentous oligomers with a helical architecture on the lipid membranes in vitro^6^. Conversely, NADP+ binding results in inactive ternary complexes that remodel membranes into spherical assemblies of branching tubes in vitro^7^, resembling the PLB ultrastructure known as a cubic phase. As NADPH can displace NADP+ in the binding pocket, forming an active complex and maintaining the cubic phase, it has been suggested that fluctuations in the NADP+:NADPH ratio play a role in PLB development in etiolated plants^7^.

LPOR originated approximately 1.5 billion years ago in cyanobacteria^8,9^, likely as an adaptation to rising atmospheric oxygen levels^10^, which hindered the activity of the alternative enzyme catalyzing the same reaction, namely dark-operative Pchlide reductase (DPOR)^11^. The two enzymes differ in reaction mechanism, electron donors, structural architecture, and phylogenetic origin, serving as examples of convergent evolution^12^. The cyanobacterial LPOR and DPOR were subsequently inherited by green algae and plants, where the sequence of the former underwent modifications that affected its lipid-regulated properties. In cyanobacteria, MGDG promotes release of the product of the reaction, chlorophyllide (Chlide), from LPOR^7^, whereas in plants, MGDG binds within the LPOR binding pocket, interacting directly with the pigment, influencing its fluorescence emission, and promoting the oligomerization^6,13^. Two regions of the protein, known as oligomerization interfaces I and II, are responsible for this process. It was suggested that insertions and deletions within the LPOR gene, which occurred in green algae, gave rise to a plant-like LPOR homolog capable of forming PLB structures^7^.

Interestingly, the ancestors of angiosperms lost the DPOR genes, making Chl biosynthesis entirely reliant on LPOR’s photocatalytic activity in all modern-day flowering plants, in contrast gymnosperms, which uses both DPOR and LPOR^14^. Over the last 200 million years, the LPOR gene has undergone duplication events independently in both monocots and eudicots, leading to the emergence of two distinct isoforms in each group^8^ (Fig. 1). In eudicots, one of these isoforms further duplicated, giving rise to a third isoform independently in species like tomato, Arabidopsis, and sunflower (Fig. 1). Some species, such as pea, lost one of these homologs and now depend on a single LPOR gene^8^. The roles of multiple isoenzymes within a single species remain unclear. While some studies suggest that the isoforms may be functionally interchangeable^15^, each isoform is expressed differently depending on the developmental stage and light conditions^2,16–19^, indicating potential specialization. Therefore, in this paper, we decided to characterize the properties of 13 isoforms of plant LPORs: from pine, wheat, sunflower, tomato, pea, and Arabidopsis.

**Figure 1.**
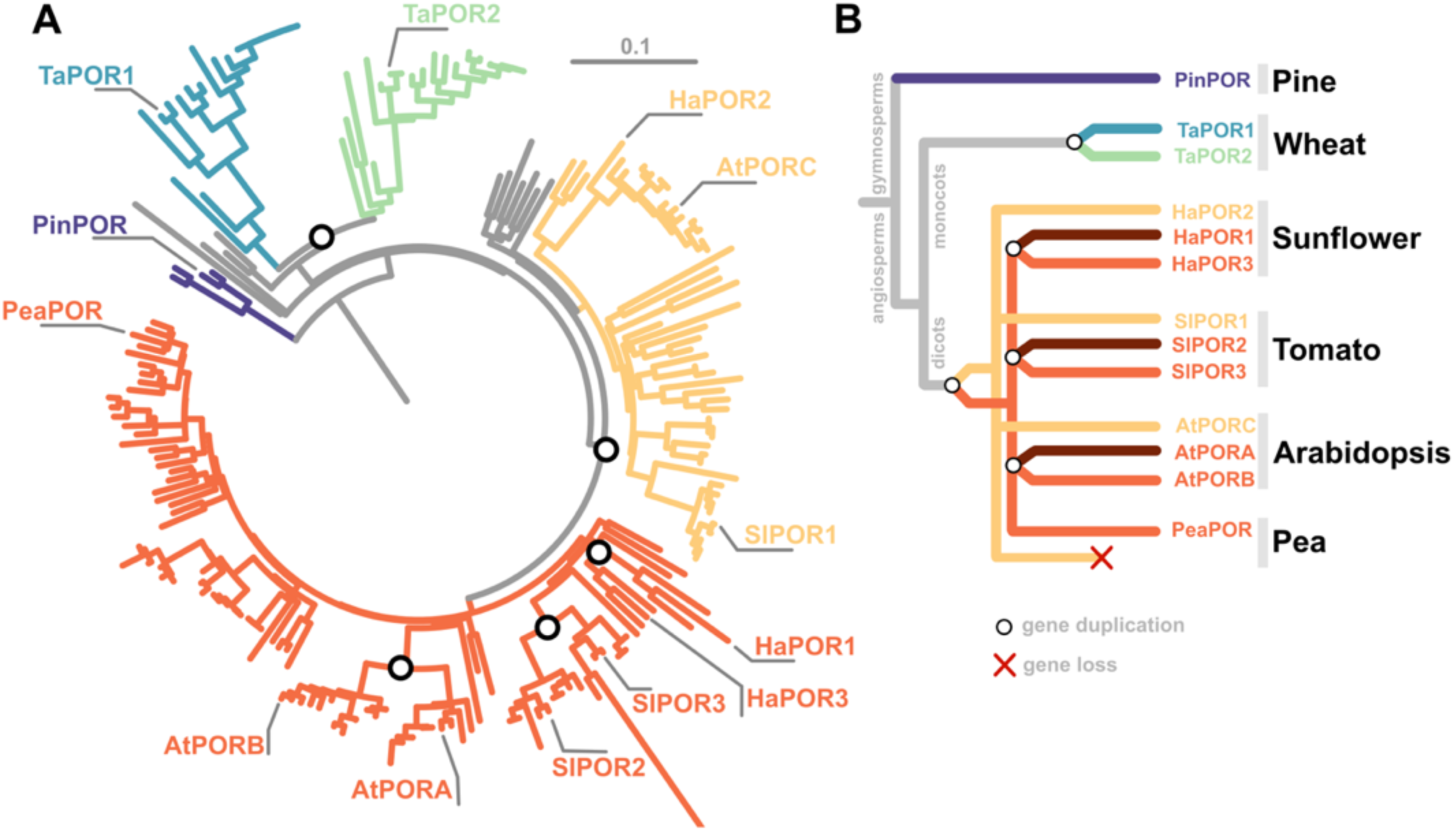
Phylogeny of LPOR isoforms in selected plant species. **A.** Rooted phylogenetic tree of seed plants. **B.** The simplified phylogenetic tree showing the origin of multiple isoforms in selected plant species.

One of the oldest methods applied to study the enzyme’s activity is fluorescence measurement at 77 K, dating back to 1957^20^. Freezing the sample inhibits the reaction, allowing for excitation of the pigment and subsequent emission measurements from the photocatalytic LPOR complexes, that would otherwise undergo photoconversion at room temperature. At the same time, low temperature makes the fluorescence peaks narrower, enabling precise tracking of even small shifts in the emission maximum. Those shifts carry the information about the pigment microenvironment, since Pchlide fluorescence emission is sensitive to the surroundings, for instance to the polarity of solvent^21^ or the local environment, such as whether the molecule is membrane-bound or soluble^22^. In PLBs, Pchlide exhibits a characteristic emission at 655 nm^23,24^, which can be reconstituted in vitro using a mixture of the pigment, NADPH, LPOR isoform from plant, and MGDG-containing lipids^13^. In this study, by utilizing low-temperature fluorescence measurements, we identified differences in Chlide binding, NADPH binding, and oligomerization within the investigated set of isoforms. With this data, along with insights from structure-predicting algorithms, we proposed a mechanism for substrate binding that leads to oligomerization.

## Results

### Composition of reaction mixtures and emission maxima

To understand the relationship between the composition of the LPOR complex and its emission maximum, we prepared a set of reaction mixtures with AtPORB (Fig 2AC) and measured low-temperature fluorescence emission spectra. We used either 5 µM Pchlide (Fig. 2A) or Chlide (Fig. 2C), with or without 200 µM NADPH or NADP+, and in lipid-free environment, or with 400 µM lipids. We used either phosphatidyl glycerol (PG) or a lipid mixture which has been previously utilized in the cryoEM study of plant AtPORB and referred to as the OPT lipids (50 mol% MGDG, 35 mol% DGDG, and 15 mol% PG).

**Figure 2.**
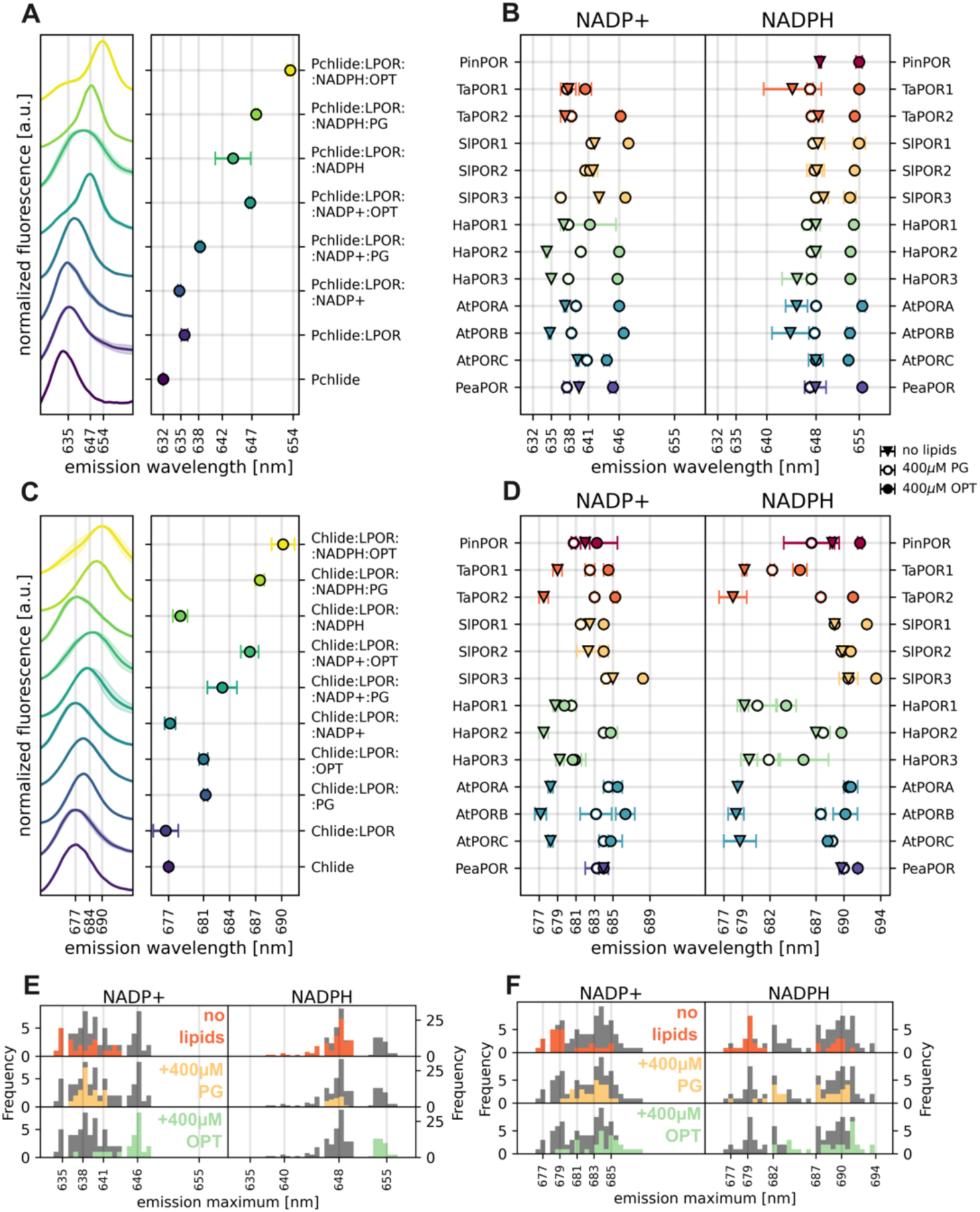
The composition of the reaction mixture affects the emission maxima of Pchlide and Chlide. **AC.** Normalized fluorescence emission spectra of reaction mixtures with Pchlide (A) and Chlide (C), along with their emission maxima. Shaded areas represent the standard deviation between replicates. **BD.** Fluorescence emission maxima of complexes with Pchlide (B) and Chlide (E) in the presence of NADP+ (left panels) or NADPH (right panels) for 13 plant LPOR isoforms, measured in a lipid-free buffer, with PG, or with OPT lipids. **EF.** Distribution of the emission maxima of Pchlide (C) and Chlide (F) shown in panels B and E. The overall distribution of data points for each pair of pigment and dinucleotide is shown in gray. **A-F**. The reaction mixtures contained 15 µM LPOR, 5 µM pigment, 200 µM dinucleotide, and 400 µM lipids in various combinations.

We observed that both pigments in complexes with AtPORB exhibited emission maxima that were generally more red-shifted in the presence of NADPH compared to NADP+ (Fig. 2AC). Similar effect for observed for samples with OPT lipids comparing to the samples that have not been supplemented with lipids. The determined shifts of emission maxima reflect the ability of AtPORB to bind the given set of a pigment and a dinucleotide, in a presence or absence of lipids. To probe the landscape of properties of other LPOR homologs we investigated the spectra of Pchlide:LPOR complexes with NADP+ and NADPH for 12 other plant isoforms, originating from 6 different species including those with only one LPOR isoform, as well as those with two or three (Fig. 1). We analyzed one isoform from pine Pinus mugo (Uniprot: Q41202; hereahter: PinPOR), two from wheat Triticum aesivum (Uniprot: CAA54042 and W4ZSC8; hereafter: TaPOR1 and TaPOR2), three from tomato Solanum lycopersicum (Uniprot: K4CFW6, K4CXM0 and K4DCQ6; hereafter: SlPOR1, SlPOR2 and SlPOR3), three from sunflower Helianthus annuus (Uniprot: OTG07258, OTG15405 and OTG26264; hereafter: HaPOR1, HaPOR2 and HaPOR3), three from Arabidopsis thaliana (Q42536, P21218, O48741; hereafter: AtPORA, AtPORB, and AtPORC) and one from pea Pisum sativum (Uniprot: CAA44786; hereafter PeaPOR).

We observed that the isoforms exhibited a range of emission maxima even for the exact same conditions (Fig. 2BD, S1). To better visualize the diversity of these emissions and the effect of added lipids, we calculated the distribution of the determined maxima (Fig. 2EF). For samples containing NADP+ and Pchlide, the emission maxima varied within an 8-nm range; however, the addition of PG narrowed this range to around 638 nm. OPT lipids shifted emissions for most isoforms to 646 nm, except for TaPOR1, SlPOR2, and HaPOR1. In contrast, reaction mixtures with NADPH and Pchlide exhibited greater uniformity. In a lipid-free environment, most isoforms emitted at approximately 648 nm, with the exception of TaPOR1, HaPOR3, AtPORA, and AtPORB. The presence of PG resulted in emissions at ∼648 nm all isoforms, whereas the addition of OPT lipids shifted the emission to around 655 nm.

Significant differences among isoforms were evident in samples with Chlide, especially those without lipids. Two groups can be distinguished: one with Chlide emission below 680 nm and the other, with emission maxima above 680 nm (Fig. 2F). In the absence of lipids, the former group likely represents unbound Chlide or LPOR:Chlide complex without any dinucleotide bound, and the latter group consists of complexes with either NADP+ or NADPH, with emissions at ∼684 nm and ∼687 nm, respectively (Fig. 2C). The isoforms that bind Chlide and NADP(H) most effectively in a lipid-free environment included PinPOR, SlPORs, and PeaPOR. Interestingly, Chlide:HaPOR2 was able to bind NADPH but not NADP+. Overall, the addition of PG and OPT lipids induced a red shift in the emission maxima of the complexes with Chlide, similarly to those with Pchlide. The effect of PG was most pronounced for TaPORs and AtPORs, while OPT lipids caused a red shift in the emissions of NADPH-containing samples for PinPOR, TaPOR2, SlPORs, HaPOR2, AtPORs, and PeaPOR (Fig. 2D).

These results indicate that all investigated plant LPOR isoforms can, in principle, form the same complexes, however, they may differ in their required concentrations of substrates and lipids. To further explore this, we examined NADPH binding and the effect of OPT lipids across a range of concentrations for all isoforms.

### NADPH binding

We analyzed the NADPH binding in unilluminated samples containing 5 µM Pchlide and 15 µM LPOR, without and with the addition of 400 µM OPT lipids. We observed that the increasing NADPH concentration in the reaction mixture led to the higher emissions at ∼648 nm for samples without OPT lipids and at ∼655 nm for those with OPT lipids. (Fig. 3A, S2). To quantify this, we calculated the relative fluorescence at either 648 or 655 nm and plotted it against the NADPH concentration (Fig. 3B, S3). The resulting plots resembled saturation curves, to which a modified Michaelis-Menten equation could be fitted (Fig. 3B, S3, Materials and Methods). From these plots, we determined the K_M_^D^ constant, which in this context represents the NADPH concentration required to increase the emission at 648 or 655 nm to half of the maximum level theoretically achievable with an infinite NADPH concentration, therefore it reflects the NADPH binding affinity.

**Figure 3.**
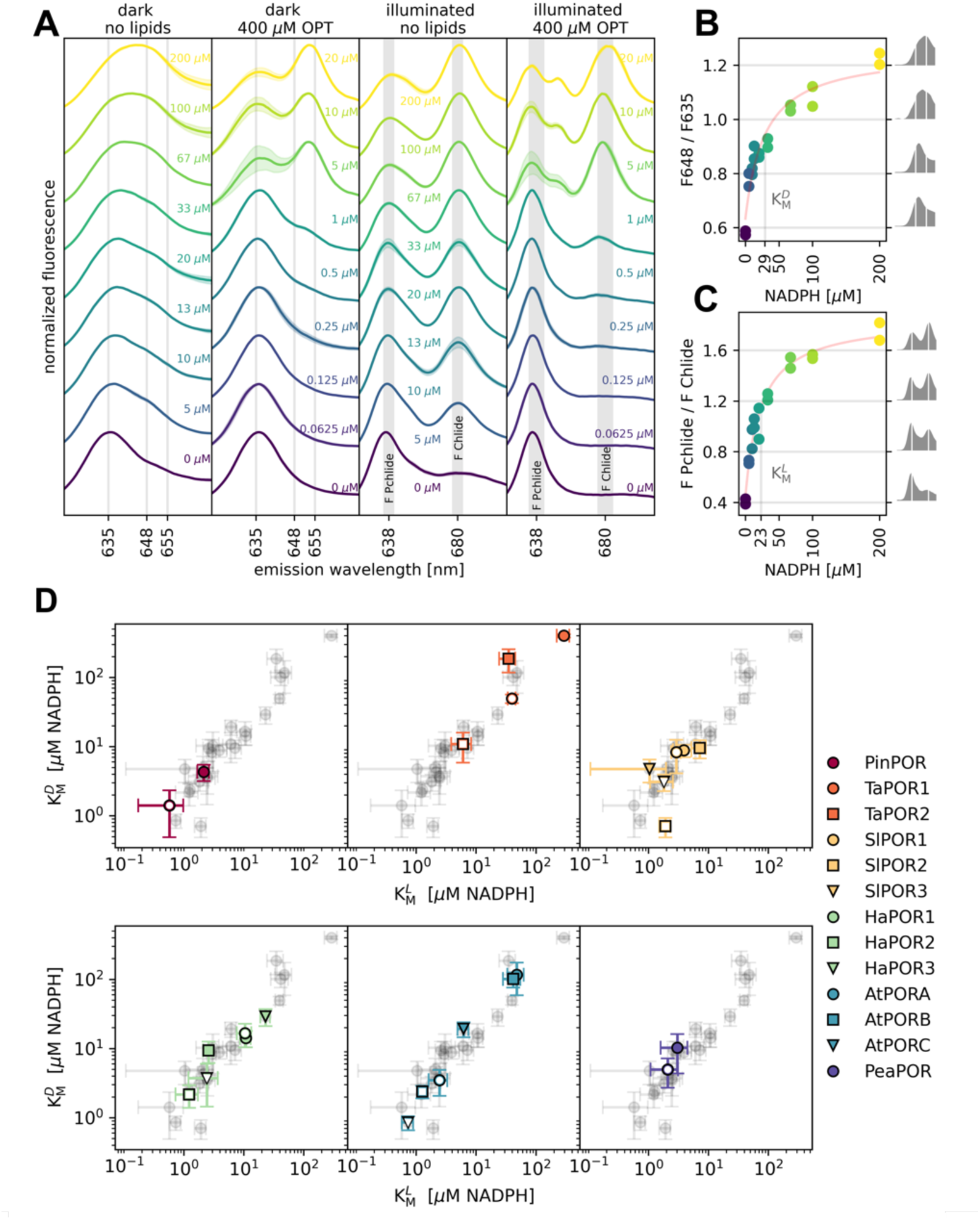
Low-temperature fluorescence emission spectra enable monitoring of NADPH binding and product formation. **A.** Normalized spectra of reaction mixtures containing 5 µM Pchlide, 15 µM HaPOR3, with or without the addition of OPT lipids, and varying NADPH concentrations (indicated next to the spectra), measured before (dark) and after illumination (illuminated). Shaded areas represent the standard deviation between replicates. **BC.** Relationship between NADPH concentration and: (B) the intensity ratio of 647/635, (C) the relative intensities of the generated product (F Chlide) and the remaining substrate (F Pchlide). A fit of a modified Michaelis-Menten equation is shown, along with the graphical representation of K_M_^D^ (B) or K_M_^L^ (C). Spectra corresponding to the given F647/F635 values (B) or F Chlide/F Pchlide values (C) are shown on the right side of the plot. **D.** Relationship between K_M_^D^ and K_M_^L^ for investigated isoforms. The overall distribution of data points is shown in gray. Error bars represent the uncertainty associated with the fitted constant K_M_^x^. The values of the constants are presented in Table S1.

All samples were thawed in darkness, illuminated for 20 seconds to initiate the reaction, and then frozen again and remeasured. As expected, increasing NADPH concentration in the reaction mixture led to greater Chlide fluorescence (Fig. 3AC, S4). We then calculated the relative Chlide fluorescence (F Chlide/F Pchlide) and plotted it against NADPH concentration (Fig. 3C, S5). These plots also resembled saturation curves, to which a modified Michaelis-Menten equation could be fitted (Fig. 3C, S5 Materials and Methods). From these, we determined the K_M_^L^ constant, representing the NADPH concentration required to produce Chlide fluorescence equal to half of the maximum achievable with infinite NADPH concentration. Thus, K_M_^L^ is a kinetic constant, while K_M_^D^ reflects NADPH binding affinity.

We observed a linear correlation between K_M_^D^ and K_M_^L^ determined for all of the investigated isoforms when plotted on a logarithmic scale (Fig. 3D), suggesting that the red-shift, which reflects NADPH bound to LPOR, correlates with higher activity. However, K_M_^L^ values were consistently lower than K_M_^D^ values. TaPOR1, TaPOR2, AtPORA, and AtPORB exhibited the weakest NADPH binding in the lipid-free environment, with K_M_^D^ constants greater than 100 µM. Conversely, PinPOR, all SlPORs, HaPOR2, and PeaPOR effectively bound NADPH in a lipid-free environment, with K_M_^D^ constants below 10 µM. The addition of 400 µM OPT lipids lowered the K_M_^D^ constant for all isoforms, except for SlPOR1, SlPOR3, HaPOR1, HaPOR2, and PeaPOR, where the constant remained unchanged within error estimates. Generally, the higher the K_M_^D^ constant in a lipid-free environment, the greater the effect of OPT lipids.

The determined distribution of K_M_ constants revealed that pine tree, pea, and tomato LPOR isoforms exhibit high activity and strong NADPH binding, in contrast to the wheat isoforms. Arabidopsis and sunflower isoforms displayed diverse properties, indicating variability in their activity and NADPH binding capabilities, at least under the conditions of the experiment.

### Effect of OPT lipids

Next, we investigated how the florescence emission spectrum of reaction mixture containing 5 µM Pchlide, 15 µM LPOR and 200 µM NAPDH changes in the presence of OPT lipids. We observed that increasing the concentration of OPT lipids in the reaction mixture initially led to a higher band at ∼655 nm, but beyond a certain concentration, a band at ∼635 nm began to appear (Fig. 4A, S6). To visualize this process, we calculated the logarithm of the intensity ratio at 658 nm to that at 631 nm and plotted it against the OPT concentration (Fig. 4B). Since the emission at ∼655 nm is associated with LPOR oligomers^6^, the log(F658/F631) can be interpreted as an index of oligomerization. Nearly all analyzed isoforms displayed a concave-down curve, except for SlPOR1 and PeaPOR, which showed a continual increase in log(F658/F631) with more lipids, at least within the tested range of OPT lipids. For all other isoforms, the optimal OPT concentration was 40-200 µM. The highest values of the oligomerization index were observed for PinPOR and HaPOR2, while TaPOR1 had the lowest value of log(F658/F631) within its optimal concentration range.

**Figure 4.**
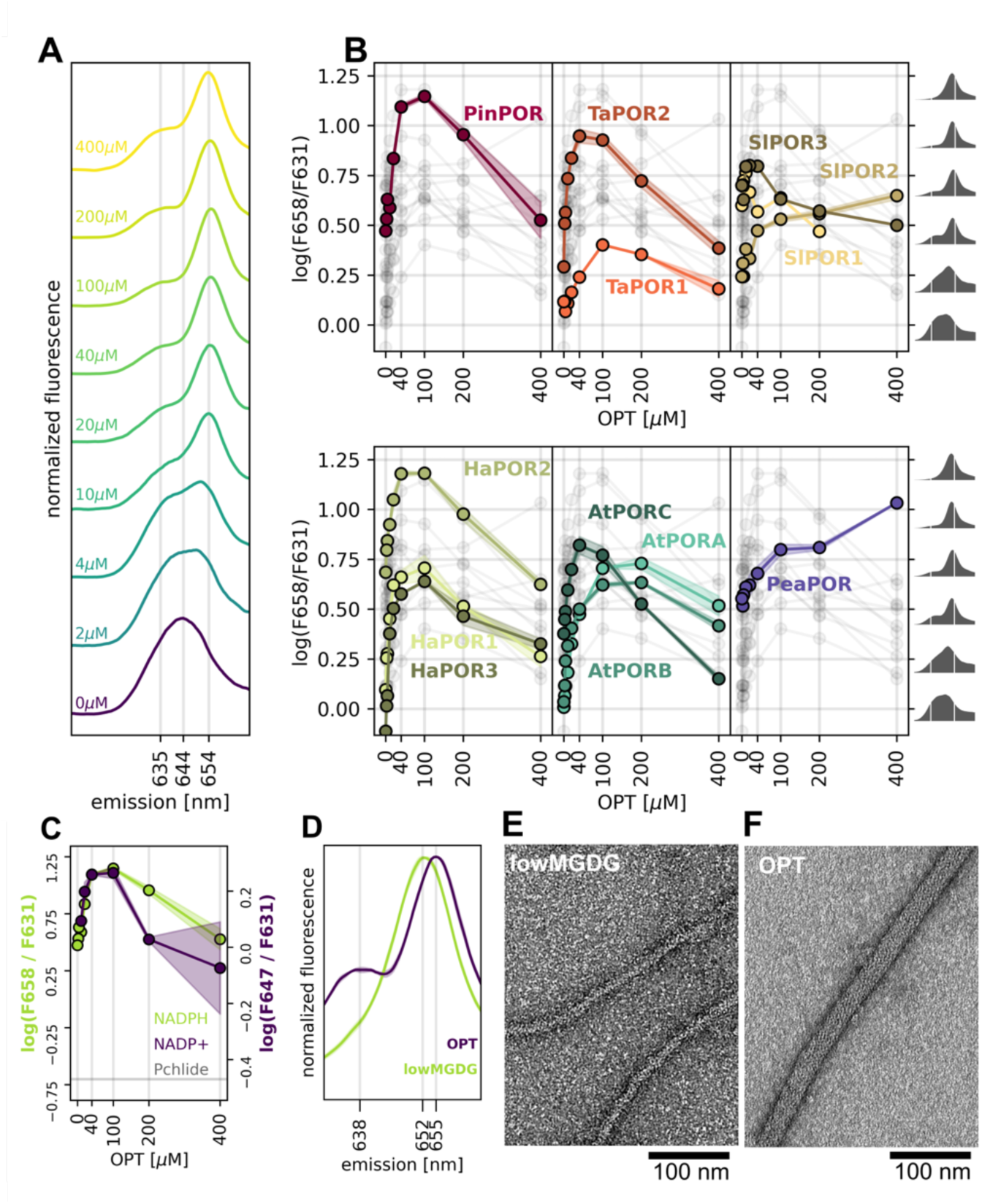
Lipids affect the spectrum of the LPOR ternary complex. **A.** Normalized spectra of reaction mixtures containing 5 µM Pchlide, 15 µM HaPOR3, 200 µM NADPH, and varying concentrations of OPT (indicated next to the spectra). **B.** Relationship between the base-10 logarithm of the intensity ratio (658/631) and OPT concentration. Spectra corresponding to the given log(F658/F631) values are shown on the right side of the plot. Shaded areas represent the standard deviation between replicates. The overall distribution of data points is shown in gray. **C.** The relationship between OPT lipid concentration and the relative fluorescence intensity of PinPOR complexes with 200 µM NADPH (655/632, green series, left axis) or with NADP+ (647/632, purple series, right axis). A gray horizontal line labeled “Pchlide” represents the 655/632 and 647/632 ratios for pure Pchlide in buffer. Shaded areas represent the standard deviation between replicates. **D.** Normalized spectra of reaction mixtures containing 5 µM Pchlide, 15 µM AtPORB, 200 µM NADPH, and either 100 µM OPT lipids or a lipid mixture with low MGDG concentration (lowMGDG: 2.5 mol% MGDG, 47.5 mol% PG, 50 mol% DGDG). **EF.** Negative-stain transmission electron microscopy micrographs of reaction mixtures containing 5 µM Pchlide, 15 µM AtPORB, 200 µM NADPH, and either 100 µM lowMGDG lipid mixture (E) or OPT lipids (F). The presented structures are typical and frequently observed for each composition of the reaction mixture.

Additionally, we investigated the effect of OPT lipids using samples supplemented with NADP+ instead of NADPH. For these samples, we measured the normalized emission at 647 nm rather than 658 nm, since Pchlide complexes with OPT lipids and NADP+ emit at 647 nm (Fig. 2A) and are known to form a cubic phase^7^. Consequently, the log(F647/F631) parameter for NADP+ samples can be interpreted as an index of oligomerization into the cubic phase. Our recent work demonstrated that for AtPORB, the relationship between OPT concentration and oligomerization indices calculated for both NADPH and NADP+ is similar^7^. As shown in Fig. 4C, this is also true for PinPOR, suggesting that it is likely applicable to other isoforms as well.

Besides affecting the oligomerization index, the OPT concentration also influenced the emission maximum of the complex. As more OPT lipids were added, the emission gradually red-shifted, reaching up to 656 nm. However, beyond a certain concentration, the emission of the main band shifted towards the blue end of the spectrum by approximately 3 nm, except for SlPORs and PeaPOR (Fig. S7). Moreover, we found that using a different lipid mixture in the reaction, specifically one with a low MGDG content (lowMGDG: 2.5 mol% MGDG, 47.5 mol% PG, 50 mol% DGDG), produces thin LPOR filaments with a diameter of ∼15 nm, compared to the ∼24 nm diameter produced with OPT lipids (Fig. 4EF). Interestingly, these complexes also differed in their emission maxima. The thin filaments emitted at 652 nm, whereas filaments produced with OPT lipids emitted at 655 nm (Fig. 4D).

These observations suggest that the ratio of LPOR to lipids, as well as the relative MGDG concentration, is crucial for the oligomerization process. The oligomers that differ in their emission maximum likely differ in their Pchlide microenvironments within the binding pocket what affects the subunit organization within the oligomer, and therefore the oligomer architecture.

### Emission maxima before and after illumination

The analysis of NADPH binding and the effect of OPT lipids yielded a dataset comprising 642 samples with varying NADPH and OPT concentrations. For each sample, we measured the emission spectrum before and after illumination. Then we selected those samples where the emission maximum of Chlide could be reliably determined, that is those with NADPH concentrations at least 1 µM. This resulted in 472 pairs of Pchlide and Chlide emission maxima, which are plotted in Fig. 5A along with their distribution.

**Figure 5.**
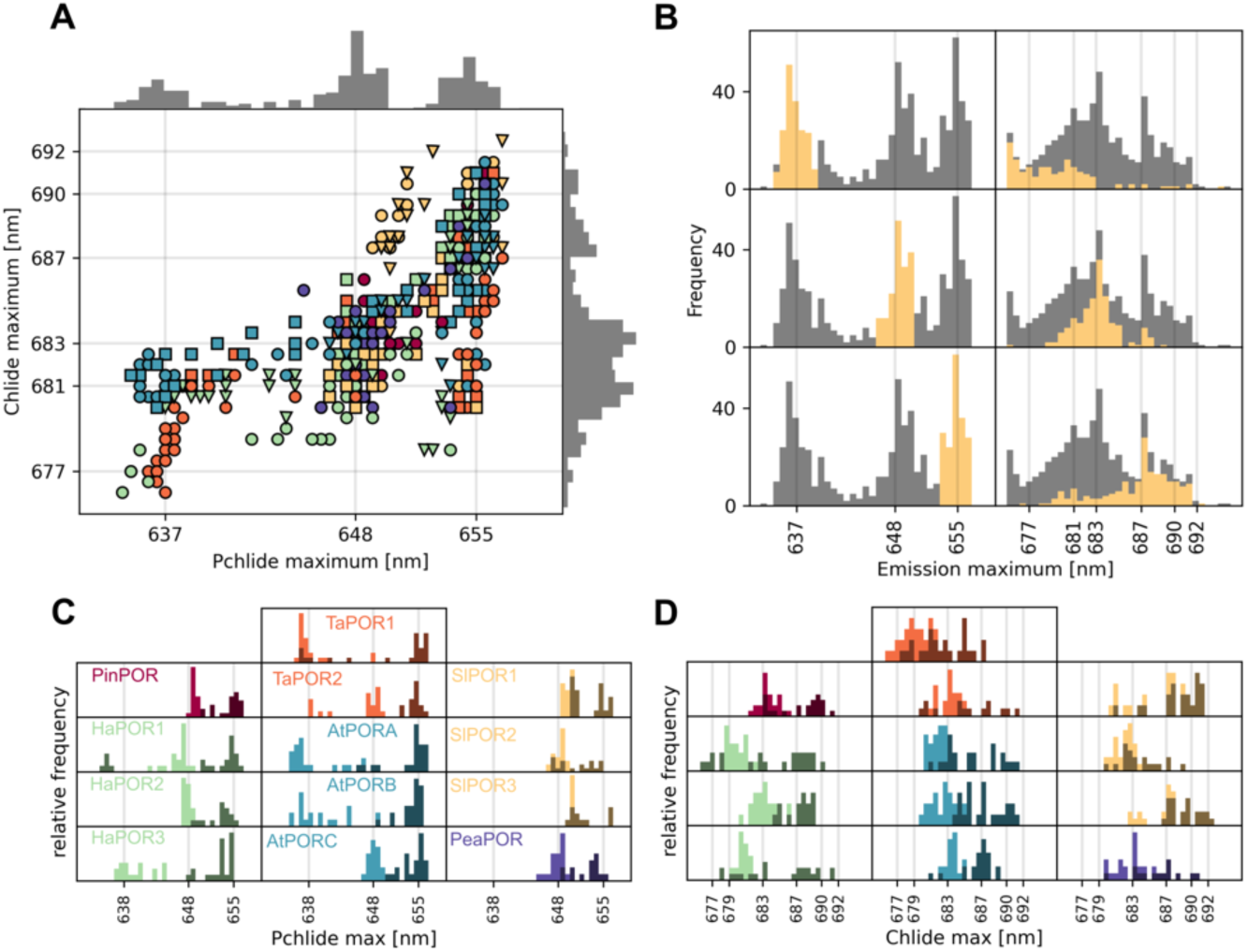
Relationship between emission maxima of Pchlide and Chlide generated after illumination of the reaction mixture. **A.** Emission maxima of samples ([NADPH] of at least 1 μM), measured before and after 20 seconds of illumination, for 13 different LPOR isoforms. Marker color and style correspond to those used in Fig. 3. The distribution of data points is shown on the top and right axes. **B.** Distribution of emission maxima for Pchlide (left panels, yellow bars) and the corresponding maxima of Chlide generated after 20 seconds of illumination (right panels, yellow bars). Three subsets of Pchlide emissions were selected based on maxima at 637, 648, and 655 nm (± 2.5 nm). The overall distribution of data points is shown in gray. **CD.** Distribution of emission maxima for Pchlide (C) and Chlide generated after 20 seconds of illumination (D), determined for the samples ([NADPH] of at least 1 μM) presented in Figs. 3 and 4, for each analyzed isoform. The arrangement of the isoforms in D is the same as in C. The maxima of the reaction mixtures containing lipids are shown in the darker shade.

This analysis revealed that Pchlide emission maxima clustered into three groups: around 637, 648, and 655 nm. In contrast, Chlide emission maxima were more broadly distributed in the range of 675-692 nm, with the most common maxima at 681, 683, and 687 nm (Fig. 5A). We found that if the Pchlide complex emitted at 637 ± 2.5 nm, the resulting Chlide complex typically had a maximum around 681 nm or at shorter wavelength (Fig. 5B). Conversely, if the emission maximum of Pchlide was at 648 ± 2.5 nm, the Chlide emission was generally around 683 nm. For Pchlide complexes emitting at 655 ± 2.5 nm, the resulting Chlide emission often shifted to 687 nm or higher after illumination.

We then classified the distribution of Pchlide and Chlide emission maxima by isoform (Fig. 5CD) to highlight differences between the analyzed LPOR variants. We found that PinPOR, all SlPORs, HaPOR2, AtPORC, and PeaPOR predominantly formed complexes with Pchlide that emitted fluorescence at around 648 and 655 nm, while the other isoforms produced more blue-shifted complexes, especially for reaction mixture lacking the lipids, indicating weaker NADPH binding and affected formation of the active complex (Fig. 5C).

Analysis of the Chlide emission maxima revealed that TaPOR1, SlPOR2, HaPOR1, and HaPOR2 most frequently produced blue-shifted Chlide complexes, suggesting these isoforms release the reaction products more quickly. Conversely, PinPOR, TaPOR2, SlPOR1, SlPOR3, HaPOR3, AtPORA, AtPORB, and PeaPOR could form Chlide complexes with emission maxima exceeding 690 nm, but only for the reaction mixtures containing lipids (Fig. 5D). Complexes exhibiting such emission are formed only with OPT lipids and NADPH (Fig. 2), suggesting that these isoforms can replace the NADP+ generated during the reaction with a new NADPH molecule and form stable, inactive complexes on the membrane.

### Structure flexibility

In order to better understand the relation between sequence, structure and function of LPOR, we employed AlphaFold2 (AF2) predictions with subsampled multiple sequence alignments (MSAs). In this approach, we used AF2 to predict a range of conformations for a given sequence, resulting in an ensemble of predictions that we then analyzed through a variety of methods. First, we sought to identify the residue ranges of highest mobility in LPOR by calculating the root mean square fluctuation of C-alpha positions (RMSF) after alignment to a reference structure, extracted from the cryo-EM structure of the LPOR oligomer (PDB: 7JK9).

We found that, although all residues showed some flexibility, four regions are significantly more mobile than the rest. These mobile elements were identified as the: Pchlide loop (230-242), oligomerization interface I (256-263), helix α10 (316-338) and oligomerization interface II (367-375) (Fig. 6). Out of these regions, only the Pchlide loop was highly conserved within plant homologs, and nearly identical among all of the 13 investigated isoforms (Fig. S8). Conversely, both oligomerization interfaces are highly diverse, while helix α10 has two variable residues at positions 325 and 338 (Fig. 6A, S8).

**Figure 6.**
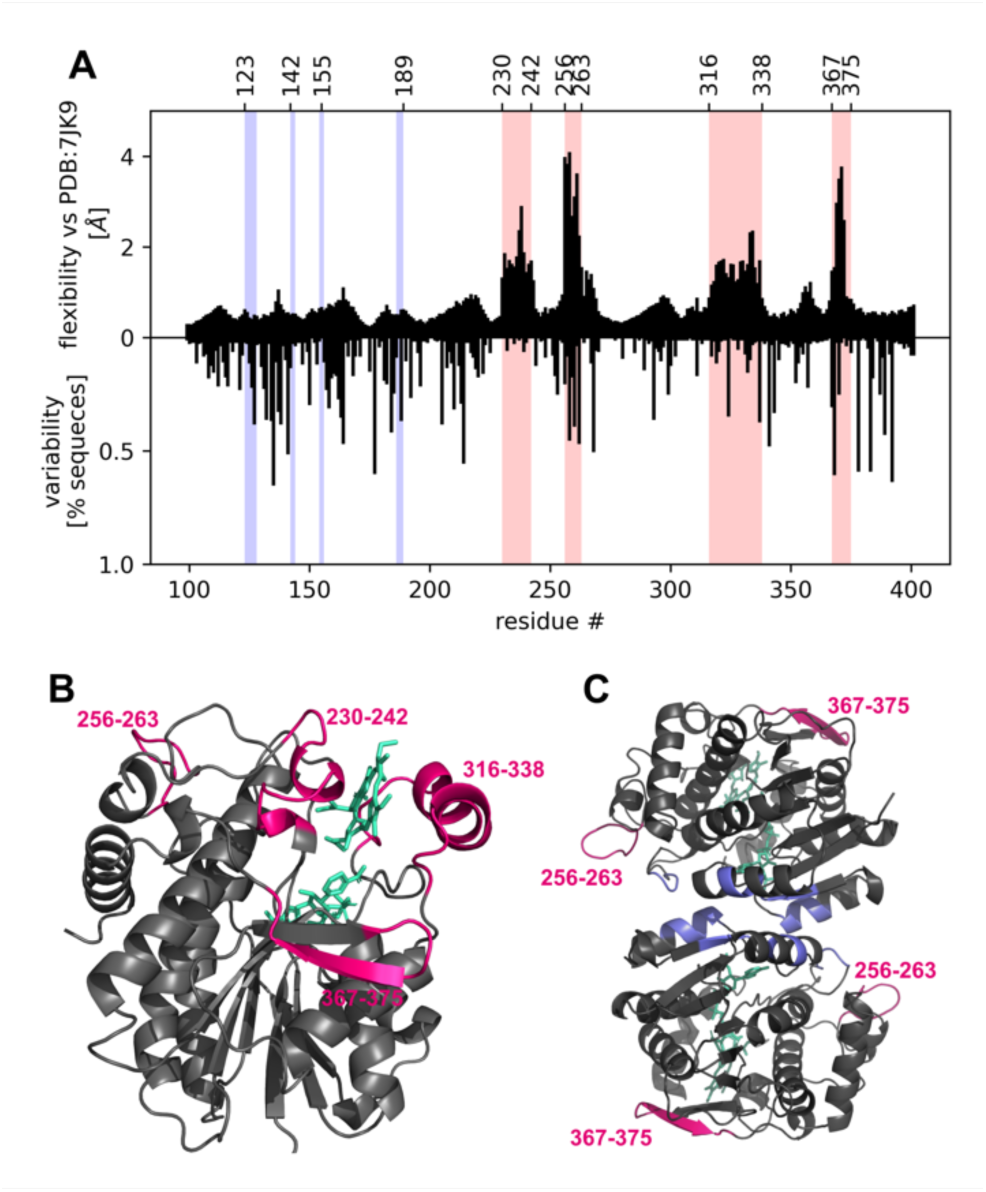
LPOR in plants has flexible regions. **A.** Root mean square fluctuation of alpha carbon atoms (flexibility) of the AtPORB predicted ensemble, calculated from 480 independent AF2 predictions, after alpha carbon alignment with the reference structure of LPOR (top panel). Variability was calculated using the previously published dataset^8^ of 194 LPOR sequences of seed plants and it represents the percentage of sequences that do not share the most common residue at a given position. The four identified flexible regions are highlighted in pink: Pchlide loop (230-242), oligomerization interface I (256-263), helix α10 (316-338), and oligomerization interface II (367-375). Residues involved in dimer formation are highlighted in blue. **BC.** Four flexible regions of LPOR (pink) highlighted on the LPOR structure of a monomer (B) and dimer (C), extracted from the cryo-EM structure of the LPOR oligomer. Residues involved in dimer formation are highlighted in blue.

We utilized AF2 to predict 480 conformations for each investigated isoform and compared these predictions to the reference structure (PDB: 7JK9) by calculating the root mean square deviation (RMSD) for the alpha carbon atoms of four identified flexible regions after alpha carbon alignment to the reference structure. A subset of these predicted conformations is illustrated in Fig. 7A, and the animated morph transitions between the predicted conformations and the reference structure are shown in Video S1. The results indicate that the Pchlide loop can uncoil and shift, the oligomerization interface I can be reoriented, and the beta sheet of oligomerization interface II can bend significantly. Additionally, helix α10 is predicted to bend and move, altering the size of the pigment-binding pocket. Analysis of the RMSD distribution reveals a bipodal character for each flexible region (Fig. 7B). Clusters that closely resemble the reference structure correspond to a single conformation, whereas clusters with higher RMSD values show greater diversity, as different conformations can exhibit similar RMSD values. Notably, the isoforms exhibit variations in the shapes of these distributions, even for regions that were nearly identical across the investigated sequences (e.g., Pchlide loop, residues 230-242, Fig. 7B). This suggests that the entire protein sequence must be considered when investigating the flexibility of specific regions. Despite the variability of the sequence of the N-terminal part (residues 100-230), it folds into a rigid structure that provides scaffolding for crucial mobile elements of the enzyme—namely, the pigment and MGDG binding pockets, as well as the two oligomerization interfaces. Although these elements are distant within the sequence, they are adjacent in the folded structure. This suggests that pigment and MGDG binding likely induces a conformational change in the binding pocket, which is then transmitted to the oligomerization interfaces.

**Figure 7.**
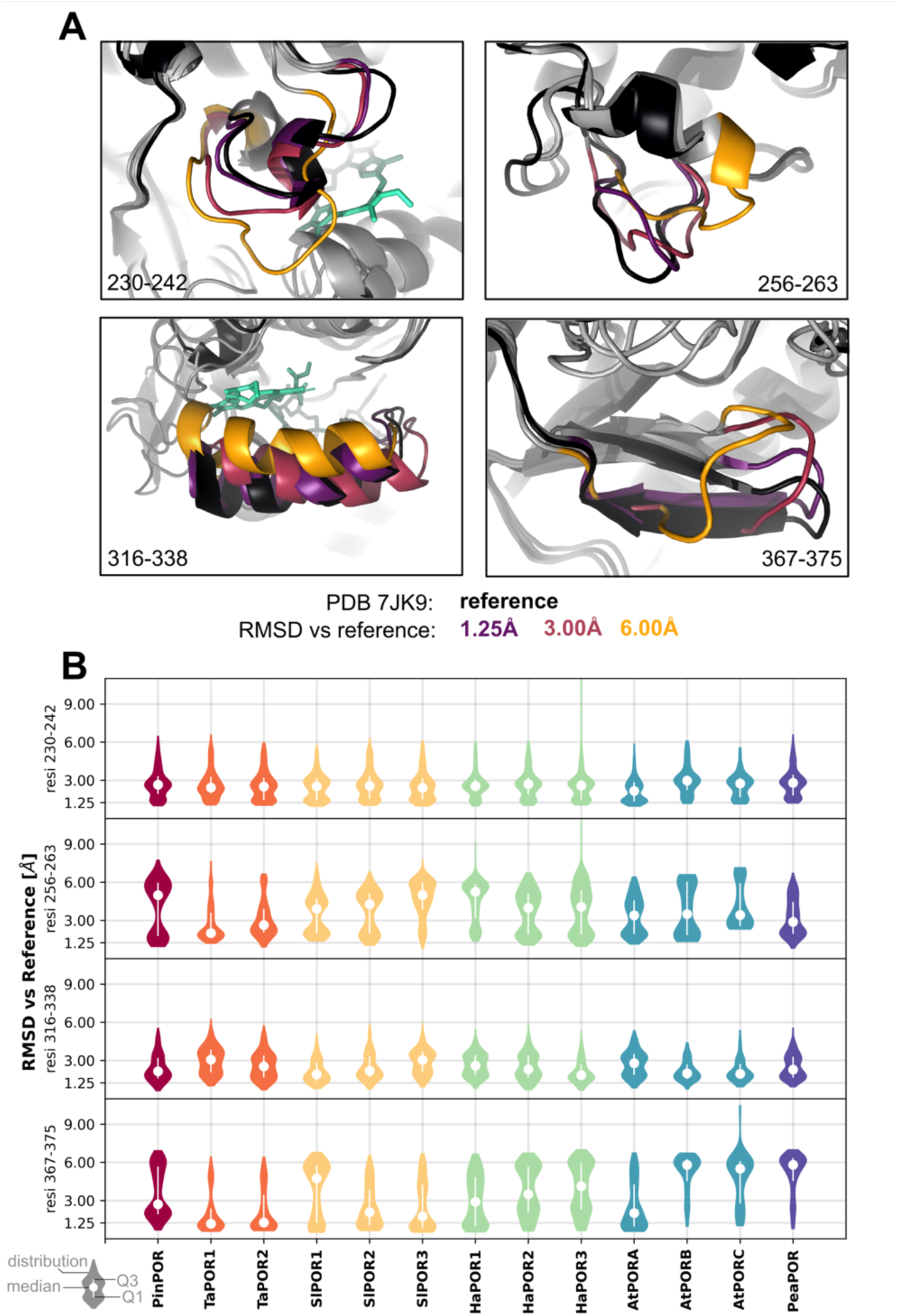
LPOR isoforms differ in the conformations that the flexible regions can adopt. **A.** Representative conformations determined for AtPORB. The reference structure (black) was extracted from the cryoEM structure of the LPOR oligomer while the alterative conformations were predicted with AF2. **B.** Distribution of root mean square deviation (RMSD) calculated for the alpha carbon atoms of four flexible regions between the predicted conformations and the reference structure. Median and the range of the first and third quadruple are marked with a white dot and white lines.

## Discussion

### Why isoforms are useful

LPOR is one of the several enzymes involved in tetrapyrrole biosynthetic pathway that has at least to two isoforms in Arabidopsis. Others include glutamyl-tRNA reductase (GluTR), aminolevulinate dehydratase (ALADH), ferrochelatase (FeCh) and ChlI subunit of magnesium chelatase. The roles and differences between these isoenzymes remain unclear, however, based on mathematical modeling of the circadian clock, it has been suggested that a given biological system becomes more stable and robust when two isoforms of the enzyme coexist and there is a significant difference in their Michaelis-Menten constants^25^. Our data indicate that for LPOR isoforms in wheat, Arabidopsis, and sunflower, there are indeed significant differences in their K_M_ constants (Fig. 2D), particularly when including values determined for lipid-free and lipid-supplemented reaction mixtures. This suggests that the role of these isoforms is to enhance the stability of Chl biosynthesis across a wide range of NADPH concentrations. On the other hand, knock-out mutants of AtPORB and AtPORC in Arabidopsis have shown that these isoforms are interchangeable and functionally redundant in mature plants^15^. However, this conclusion was derived under controlled laboratory conditions, not in real-world fluctuating environments where differences between knock-out and wild-type plants may emerge. Surprisingly, AtPORA knock-out turned out to display severe photoautotrophic growth defects and reduced apical dominance when grown under a normal photoperiod^26^, even though AtPORA is expressed only during etiolation^19^. This suggests that LPOR has functions beyond merely catalyzing Pchlide reduction. In fact, the enzyme is regarded as a regulatory hub of the Chl biosynthetic pathway for several reasons. Firstly, as a photoenzyme, its activity is influenced by external conditions, allowing it to function similarly to a photoreceptor^27^. Moreover, LPOR is known to interact with multiple enzymes and proteins involved in tetrapyrroles biosynthesis (FLU^28^, YCF54 and Chl27^28,29^, Lil3 and ChlP^30^, FeCh2^31^), while its stability depends on thioredoxins^32^ and chaperon proteins^33^. Additionally, LPOR can oligomerize into helical assembly triggered by MGDG binding^6^ and in consequence remodel lipid membranes into cubic phase^3,7^, but so far the effect of LPOR oligomeric states on the partner proteins have not been investigated. It is possible that different LPOR isoforms vary in their binding affinities for partner proteins, which could lead to distinct functional outcomes through these interactions. By modulating the expression of various isoforms, plants may fine-tune Chl biosynthesis, switching between different modes of regulation depending on environmental conditions or developmental needs, increasing their resilience or enabling them to conquer novel environmental niches. In fact, whole-genome duplications, which give rise to new isoforms and proteins, correlate with periods of extinction or global change^34^. In such harsh or disturbed environments, polyploids often thrive since the duplicated genetic material allows evolution to explore more diverse options^34^. This raises the question: what specific properties of LPOR have been favored by evolution?

### Isoforms within the same phylogenetic clades show variation

Biochemical characterization of LPOR isoforms in vitro presents significant challenges; nonetheless, several studies in the literature have attempted to tackle this issue. Isoforms from barley have been shown to differ in their Pchlide binding constants, catalytic efficacy, and the temperature dependence of NADP⁺ displacement by NADPH^35^. Our group recently reported differences in anionic-lipid-assisted NADPH binding between isoenzymes of Arabidopsis and of wheat^7,13^. Other group measured kinetic parameters of the photocatalytic reaction of selected plant LPORs revealing some differences^36^, while the others demonstrated variation in optimal pH and temperature^37^. In this study, we employed low-temperature fluorescence measurements to comprehensively characterize all isoforms across selected six plant species with diverse phylogenetic backgrounds, including both model organisms and species of agricultural importance.

The characterized isoforms belong to five different clades, which we have designated as follows: G (representing Gymnosperms), M1 and M2 (clades containing two types of isoforms from Monocots, respectively), and clades C and AB (clades containing two types of isoforms from eudicots, to which AtPORC and AtPORA/B belong to, respectively)^7^ (Fig. S9DE). Among the characterized isoenzymes, one protein belonged to clade G, one to clade M1, one to clade M2, three to clade C, and seven to clade AB (Fig. S9E).

The examined proteins in clades G, M2, and C demonstrated effective oligomerization and had a high affinity for NADPH in a lipid-free environment, with the exception of M2 (Fig. S9A-C). M1 was similar to clade AB; however, the latter exhibited the greatest diversity in terms of properties. This may be attributed to the fact that it is the largest clade in our protein set, but notably, within this clade, there were three additional duplication events that gave rise to six sub-populations of sequences (Fig. 1)^8^. The observed diversity, particularly within the clade AB, indicates that isoforms that belong to the same clade may have develop distinct functions in different plant species. One way to test this hypothesis is to identify the isoforms predominantly expressed during etiolation in different species and verify if they belong to the same clade; however, this falls outside the scope of this study and requires a dedicated investigation. So far, it was shown that LPOR mRNA level and the activity of the enzyme decreases during greening in pea, tomato and sunflower^38^.

### Diversity Among Plant LPOR Isoforms

The data presented in this study allow us to identify notable differences between the investigated isoforms. To our surprise, we observed diversity in Chlide binding, with certain isoforms—such as PinPOR, PeaPOR, and all SlPORs—showing an exceptional affinity for Chlide, particularly in the presence of lipids and NADPH. This suggests that these isoforms can retain Chlide after the reaction, especially when NADPH concentrations are high enough to displace the NADP⁺ formed within the enzyme’s binding pocket during the photocatalytic process. In contrast, reaction mixtures containing Chlide for isoforms such as TaPOR1, HaPOR1, and HaPOR3 tend to display more blue-shifted emission maxima, suggesting weaker binding. These findings suggest that LPOR isoforms differ in their rates of Chlide release and complex disassembly, potentially influencing the overall rate of the Chl biosynthetic pathway, even in mature leaves.

Another identified difference between the isoforms was the K_M_ constants measured in a lipid-free environment (Fig. 3D). Plants with only one LPOR isoform, such as pine and pea, possess isoenzymes that tend to form active complexes even at relatively low NADPH concentrations, similar to all SlPORs. In contrast, LPOR variants from sunflower and Arabidopsis exhibit diverse NADPH binding properties (Fig. 3D). Recently, we identified four residues at positions 122, 178, 312, and 318 that can affect the K_M_ constant by stabilizing the closed, active conformation of the enzyme^7^. However, it appears that predicting the K_M_ constant of an isoform based solely on these four positions is not possible, as LPOR isoenzymes with identical residues at these positions can have strikingly different K_M_ constants (for example, TaPOR2 and SlPOR3; see Fig. S10). Most likely, other positions contribute to NADPH binding and influence this process.

The isoforms also exhibited differences in their oligomerization properties. We observed that, for most isoenzymes, high lipid concentrations were associated with decreased emissions from the oligomeric form at ∼655 nm (Fig. 4B). We speculate that this phenomenon is influenced by the strength of interactions between the subunits within the oligomer. As lipid concentration increases in the reaction mixture, the density of subunits on the membranes decreases, making it more difficult to maintain the oligomeric form. The sequences of oligomerization interfaces are diverse within the analyzed subset of isoforms; therefore, it is highly probable that some interactions are stronger than others. While it is beyond our scope to determine which isoenzymes exhibit stronger interactions based on sequences alone, the data (Fig. 4B, S7) suggest that interactions between subunits within oligomers of SlPORs are weak, whereas interactions among the subunits of PinPOR, AtPORA, and both TaPORs are stronger, despite the latter’s weak affinity for NADPH. AF2 predictions also indicate variation in the conformations of the oligomerization interfaces that these isoforms tend to obtain (Fig. 7B), what also implies differences in the oligomerization properties.

The high lipids concentrations were also associated with a blue shift of the ∼655 nm band for most of the isoforms. This suggests that the architecture of the helical assembly also changes, since the variations in Pchlide microenvironment is reflected in the emission maximum. We have shown that LPOR can form diverse oligomers in terms of diameter and morphology. The diameter is affected by the level of MGDG in the lipid mixture (Fig. 4EF) and the protein-to-lipid ratio^6^, while the morphology is influenced by the dinucleotide: NADP+ induces the formation of branched tubes arranged in a cubic phase, whereas NADPH leads to the formation of linear filaments^7^. Certainly, high-resolution structures of other LPOR assemblies are needed to gain a better understanding of the oligomerization mechanism and to explain how lipid concentration influences this process. Nevertheless, based on the ensemble of AF2 predictions and experimental data, we can now speculate how the binding of the substrates can affect oligomerization.

### Oligomerization mechanism

AF2 predictions suggest that most of the LPOR protein is rigid, with four flexible regions: two involved in substrates binding (helix α10 and Pchlide loop), and two involved in oligomerization (oligomerization interfaces I and II). However, there is another oligomerization interface leading to the formation of antiparallel dimer (Fig. 6C), which is the core building block of the filamentous LPOR oligomer^6^. The residues involved in this interaction are both conserved and rigid (Fig. 6A), what suggest that LPOR may form a dimer before it binds the substrates. In fact, when the LPOR isoforms from barley were analyzed using gel filtration, their elution time indicated a mass consistent with that of dimers^39^. Other researchers proposed putative dimer architectures^40,41^, but they were inconsistent with the cryoEM structure published later^6^. It was also suggested that cyanobacterial LPOR works as a dimer^42^, however, plant and cyanobacterial homologs have different properties regarding regulation and oligoemrization^7^.

Based on our recent analysis of AtPORB mutants, we suggested that two different conformations of helix α10 can bind Pchlide: one in the apo state and the other in the active holo state^43^. Predictions from AF2, presented in this study, support this hypothesis by showing that helix α10 can bend and move. Based on the results presented in this study, we propose that the initial step in the oligomerization mechanism of LPOR involves Pchlide binding to the apo conformation of helix α10 (Fig. 8A). If NADPH is also bound, the interaction between the phosphate group of NADPH and R317 reorients the helix α10, reducing the size of the slit in the pigment binding pocket^7^, and allowing the Pchlide loop to interact with the pigment. This interaction remodels the loop, leading to the formation of the active complex. If this process occurs in darkness, the complex can interact with the membrane, facilitating the binding of MGDG. QM/MM calculations indicate that MGDG strongly interacts with both Pchlide and specific protein residues, particularly Y177 and K277^43^. Other residues, not included in the QM/MM study, such as those on the Pchlide loop (P241 and P242), likely also contribute to MGDG binding due to their proximity. We propose that the interactions between the residues, Pchlide and MGDG induce a conformational change in the Pchlide loop, and this remodeling is transmitted to the oligomerization interfaces (Fig. 8BC). Oligomerization interface I undergoes a conformational change due to the pulling force generated when the Pchlide loop interacts with MGDG (Fig. 8B), while oligomerization interface II adapts to a different conformation since the Pchlide loop is engaged with the pigment (Fig. 8C). These processes result in LPOR subunits adopting conformations at the oligomerization interfaces that promote strong interactions between the dimers, ultimately leading to the formation of strands of dimers that can assemble into a helical structure^6^.

**Figure 8.**
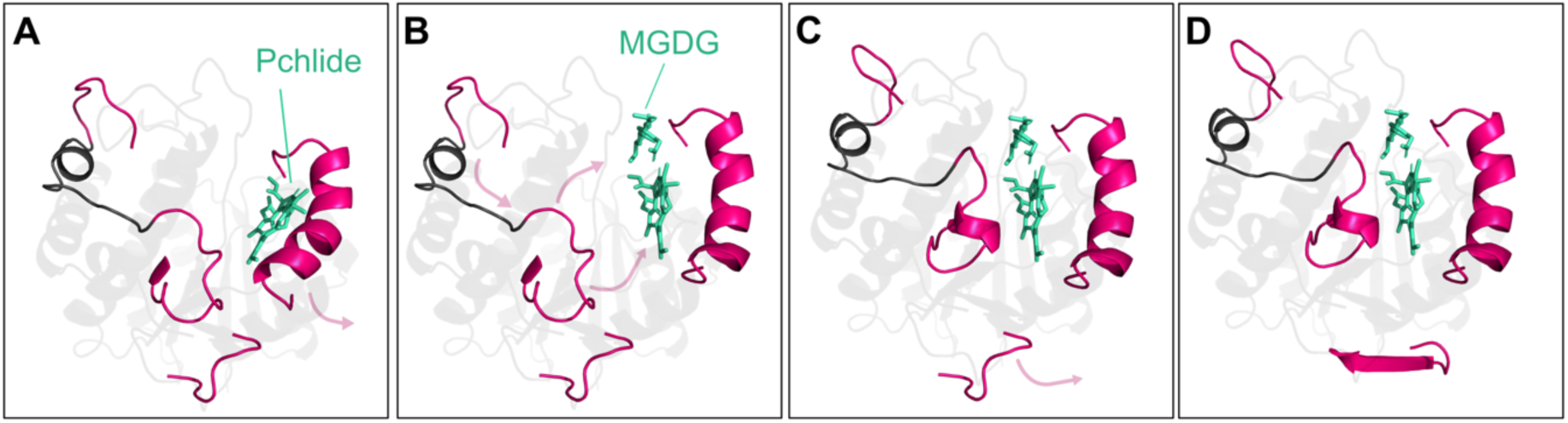
The model of conformational changes leading to the oligomerization. See the main text for a detailed explanation.

This model does not explain how NADP+ binding to LPOR:Pchlide complex can remodel lipids into the cubic phase yet. It is possible that NADP+ stabilizes an alternative conformation of the Pchlide loop and helix α10, thereby affecting the oligomerization interfaces and potentially leading to the formation of an unknown assembly that could support a cubic phase. There are organic compounds known to produce a cubic phase from helical networks^44^. Although these compounds are not biological systems, they demonstrate that helical assemblies can support cubic phase. Therefore, a dedicated topological investigation is needed to determine which architecture of LPOR subunits could potentially stabilize the ultrastructure of the PLB. Some hints into this arrangement can be inferred from the shifts of the low-temperature fluorescence emission maxima.

### Pchlide fluorescence red-shifts and the oligomeric states

The interpretation of the shifts of the fluorescence emission is a challenging issue for serval reasons. First, it has been shown that complexes of different composition can have similar emission maxima^7^. Moreover, many factors influence the fluorescence red-shift of the pigment within binding pocket of LPOR: coordination of magnesium ion within the tetrapyrrole ring^45^, interactions between enzyme’s residues and carbonyl of Pchlide^43,45^, interactions between the pigment and MGDG^13,43^, the architecture of the oligomer (Fig. 4D-F) and the redox state of the dinucleotide (Fig. 2). The latter was originally determined using homogenates of etiolated cucumber cotyledons^46^ and the emission maxima of the complexes identified in the present study closely match those reported previously^46^.

Interestingly, the addition of MGDG-containing lipids induced both the oligomerization and the fluorescence red-shift of Pchlide complexes for both NADP+ and NADPH. However, a similar red shift was also observed in lipid-supplemented Chlide complexes, at least for some of the investigated isoforms (Fig. 2). This suggests that LPOR may also form oligomers with Chlide, although this must be experimentally validated.

Notably, the red shift of Pchlide’s emission maximum upon pigment binding has been associated with strong hydrogen bonds to the keto group at C13 of the pigment^43,45^. Since complexes with NADPH exhibit a greater red shift compared to those with NADP+ (Fig. 2), it is possible that different sets of hydrogen bonds to the C13 carbonyl group are present in NADP+ and NADPH-bound complexes. As the dinucleotides do not directly interact with this region of the pigment, it is reasonable to assume that NADP+ binding induces a different conformation of the pigment-binding pocket compared to NADPH binding—most likely involving helix α10 rather than the Pchlide loop. This is because helix α10 and its downstream residues interact with both the pigment’s keto group and the nicotinamide portion of NADP(H).

Additionally, since NADPH can displace NADP+ in the binding pocket, it is possible that NADP+ is more exposed to the environment in the complex due to different conformation of the binding pocket. In the structure of the active ternary complex, NADPH is buried deep within the protein^6^. As for NADP+, although the crystal structure of a cyanobacterial LPOR homolog bound to NADP+ has been determined^47^, key regions of the enzyme—specifically the Pchlide loop, helix α10, and the downstream residues—remain unresolved. Of these regions, only the structural rearrangement of helix α10 and the downstream residues are likely to influence the exposure of the dinucleotide to the environment.

Based on these observations—specifically the interactions with the keto group at C13 and the nicotinamide moiety of NADP(H)—we hypothesize that NADP+ binding affects the conformation of helix α10, which is then translated to the oligomerization interfaces, likely through rearrangements of the Pchlide loop. However, this hypothesis requires experimental verification.

## Methods

### Gene cloning

The LPOR genes from Arabidopsis and wheat has been cloned for the previous studies^7,48^. PinPOR gene (Uniprot: Q41202), SlPOR1-3 genes (Uniprot: OTG07258, OTG15405 and OTG26264), HaPOR1-3 genes (Uniprot: OTG07258, OTG15405 and OTG26264), and PeaPOR gene (Uniprot: CAA44786) were cloned from pine, tomato, sunflower and pea, respectively with RNA extraction kit (Sigma-Aldrich) followed by cDNA synthesis with oligoT primers (ProtoScript II, New England Biolabs). The gene fragments lacking the part coding a transit peptide and pET15b plasmid (Novagen) were amplified with PCR using Q5 polymerase (New England Biolabs) according to the manufacturer’s protocol: 98 ◦C for 30 s; 30 cycles of 98 ◦C for 10 s, annealing for 30 s (see Table S2), 72 ◦C for 40 s; final extension 72 ◦C for 2 min, with specific primers (Table S2). The products of the reactions were purified from agarose gel using Gel Extraction Minipreps Kit (Bio Basic Canada) and ligated together with NEBuilder HiFi DNA Assembly Kit (New England Biolabs). Escherichia coli DH5α competent cells were transformed with the ligation mixtures and the cells containing assembled plasmid were selected on an agar medium supplemented with 100 mg/l ampicillin. The clones containing inserts were identified using colony PCR. The recombinant plasmids were isolated using plasmid purification kit (Bio Basic Canada) and the inserts were verified by sequencing (Genomed, Poland).

### Protein expression and purification

All proteins were expressed and purified according to the previously published protocol^7^ using E. coli BL21 (DE3)pRIL cells.

### Pchlide and Chlide purification

Pchlide was purified from etiolated wheat seedlings according to previous study^13^. Chlide was prepared from chlorophyll a according to the protocol^49^.

### Sample preparation and fluorescence measurements

Sample preparation was preformed according to the previous reports^7,13^. All reaction mixtures contained 37 mM sodium phosphate buffer (Na2HPO4/NaH2PO4; pH 7.1), 225 mM NaCl, 150 mM imidazole, 5 mM β-mercaptoethanol and 25% glycerol. The exact concentrations of the reagents are given in the descriptions of the figures. The range of the concentrations used in the experiments was 62.5 nM–200 μM for NADPH/NADP+ (AppliChem), 5 mM for Pchlide/Chlide, 15 μM for LPORs and 2–400 μM for the lipids (Avanti Polar Lipids). OPT lipids refer to the mixture of lipids optimized for previous structural study and contained: 50 mol% MGDG, 35 mol% DGDG and 15 mol% PG (Avanti Polar Lipids). Samples of the proteins were stored at − 20 ◦C and were gently thawed on ice just before the preparation of reaction mixtures. The addition of the pigments to the reaction mixture was performed under a dim green light. The samples were incubated in darkness for 30 min at room temperature before the measurement, then placed in quartz capillaries and frozen in liquid nitrogen for fluorescence measurement at − 196 ◦C. After the measurements, the samples were thawed at room temperature under a dim green light and illuminated for 20 s with white light of intensity 80 μmol photons m− 2 s − 1, then frozen again and measured again. Each composition of a reaction mixture for every protein had two. The low-temperature fluorescence spectra were measured with PerkinElmer LS-50 B spectrofluorometer equipped with a sample holder cooled with liquid nitrogen. The spectra were recorded at 77 K between 600 and 790 nm with a scanning speed of 400–500 nm/min. The data collection frequency was 0.5 nm and the excitation wavelength was 440 nm. Excitation and emission slits were in a range 7–10 nm. All of the spectra were normalized at their maxima.

### Calculations for K_M_

The samples were prepared as described above, and low temperature fluorescence spectrum was measured twice for each sample: before and after illumination for 20 s. From the spectrum before the illumination, the following intensity ratios were calculated: 648/635 for samples without OPT lipids, 658/635 for samples with OPT lipids. From the spectrum after the illumination the emission intensities of Chlide (F Chlide) and Pchlide (F Pchlide) were recorded at the peaks’ maxima: between 674-695 and 632–655, respectively (Fig. 3A). The ratio of F Chlide/F Pchlide was then calculated for each sample. Then, the relationship between the calculated ratios and NADPH concentrations were plotted. For every analyzed protein, a modified Michaelis-Menten equation was fitted to the data: (vmax * [NADPH])/(KM * [NADPH]) + a. The modification lies in the “+ a” term, as for inactive samples, the ratios F Chlide/F Pchlide, 648/635 and 658/635 never reach zero.

### Electron microscopy

Samples for electron microscopy were negatively stained and visulized according to the previously published protocol^7^. The presented structures were typical and frequently observed for each investigated composition of the reaction mixture.

### Phylogenetic data

The phylogenetic data used in some figures of this paper, namely the phylogenetic tree and the aligned sequences, were published as a supplementary material in our previous study^8^. iTOL^50^ was used to visualize and color the tree.

### Multiple Sequence Alignment Generation

The multiple sequence alignment (MSA) for each predicted LPOR homolog or mutant was generated using jackhmmer^51^ by querying UniRef90, small-BFD, and mgnify^52–54^, with standard parameters from the original AlphaFold2 protocol^55^ for match cut-off and thresholds.

Each MSA used for generating the predictions is included the project data directory.

### Subsampled AlphaFold2 Predictions

Conformational ensembles for POR homologs and mutants were predicted using the colabfold_batch implementation of AlphaFold2^56^ with subsampled MSAs, following the protocol described in^57^.

After running a total of 500 initial predictions with different subsampling values, we selected ‘max_seq’ and ‘extra_seq’ values of 64 and 128, respectively, as subsampling values for the following predictions.

These parameters were chosen as they led to predicted ensembles with backbone conformations similar to 7JK9 (no partially unfolded structures in the ensemble) and correctly captured the mobility of flexible residue ranges (see Table S3 for range definition).

After selecting the MSA subsampling values, we used the colabfold_batch implementation of AlphaFold2^56^ to generate 160 predictions each for the wild-type and mutant forms of AtPORB and AtPORC.

Beyond enabling dropouts and changing the MSA subsampling parameters to 64 (max_seq) and 128 (extra_seq), all other AlphaFold2 parameters were kept at their default values (5 models per prediction, 3 recycles, no templates)^56^.

Ensemble analysis of AtPORB and AtPORC and their respective mutants revealed that subsampled AlphaFold2 correctly predicted changes in ensembles due to sequence variations. We expanded our analysis to other POR homologs (See Supplemetary Table X for a list) and increased our sample size to 480 predictions for each.

### Ensemble Analysis

Calculations of alpha carbon Root Mean Square Fluctuations (RMSF) and Root Mean Square Deviations (RMSD) of atomic positions of predictions in each ensemble were conducted using MDAnalysis^58^.

RMSFs were calculated by aligning each prediction in a set to the 7JK9z reference and measuring the fluctuations of each alpha carbon within the predicted ensemble. RMSD calculations for different residue ranges were conducted by aligning each prediction in a set to the 7JK9z reference and measuring the RMSD of alpha carbon atoms in that residue range versus those in the reference.

For some predictions, such as AtPORC, re-indexing of residues in 7JK9z was necessary for accurate comparison of residue ranges. This reindexing was achieved through PyMol^59^.

For more information about the residue ranges compared, including their indexes and sequences, see Supplementary Table X.

### Representative Structure Selection

To select representative structures from each ensemble, we performed a dynamic RMSD clustering analysis with a cut-off of 3Å for selecting a new cluster centroid from each ensemble (for generating the AtPORB movies, this cut-off was 2Å).

This involved, for each prediction, calculating the alpha carbon RMSD of residues in all four residue ranges versus the positions of these same alpha carbons within the top-ranked prediction in each ensemble after aligning to it.

If that RMSD value exceeded the threshold, the prediction was appended to the list of references (becoming a cluster centroid), and all subsequent predictions were compared to it as well. Through this iterative process, we identified diverse POR conformations for each ensemble.

An abstract, system-agnostic workflow to achieve the dynamic clustering described above is available at https://github.com/GMdSilva/dynamic_structure_clustering.

Each set of representative structures is included in the project data folder.

## Acknowledgments

We thank Jerzy Kruk for his help with Pchlide purification and Leszek Fiedor for providing a sample of purified Chlide. This work was supported by SONATA project (2019/35/D/NZ1/00295) granted by National Science Centre (NCN) to MG.

