## Supplementary material for "Mobility of Four Structural Regions Drives Isoform-Specific Properties of Plant LPOR": Fig. S

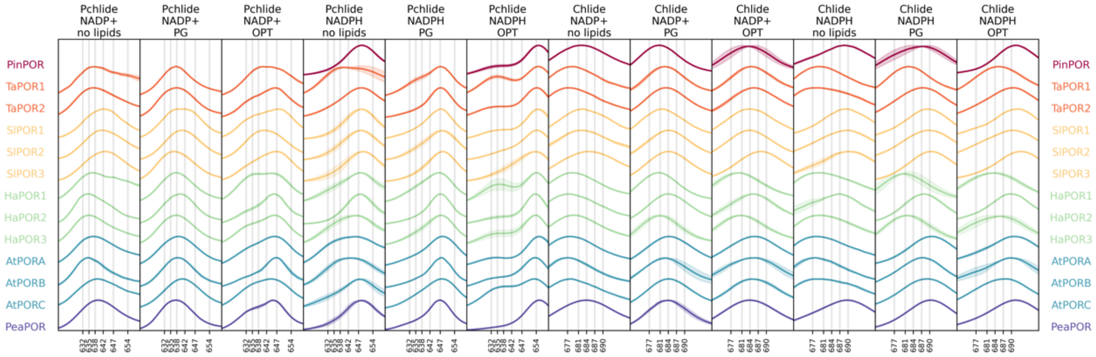


**Figure S1.** The emission maxima of reaction mixtures in different conditions for different isoforms. The reaction mixtures contained 15 µM LPOR, 5 µM pigment, 200 µM dinucleotide, and 400 µM lipids in various combinations.

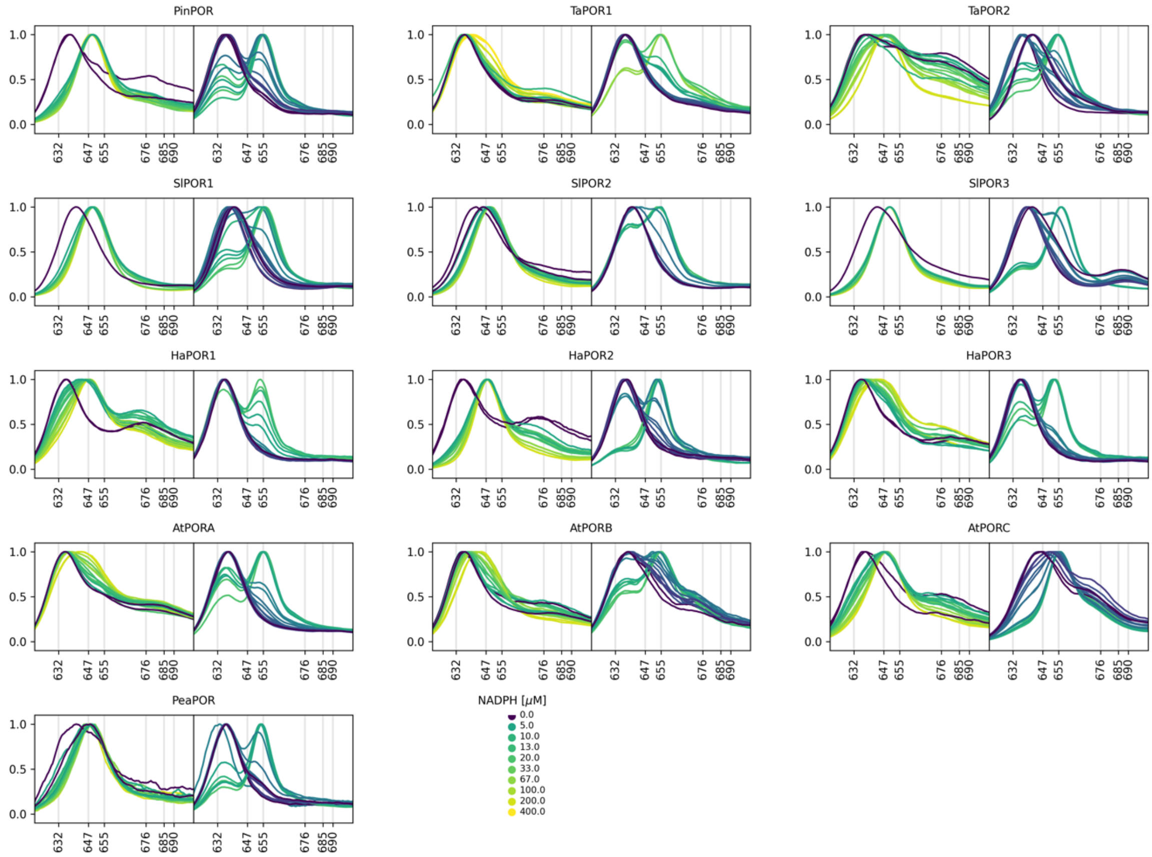


**Figure S2.** The spectra before illumination of reaction mixtures for $K_M^D$ determination without (left panels) and with 400 µM OPT (right panels) for different isoforms. The reaction mixtures contained 15 µM LPOR, 5 µM pigment and variable NADPH concentrations.

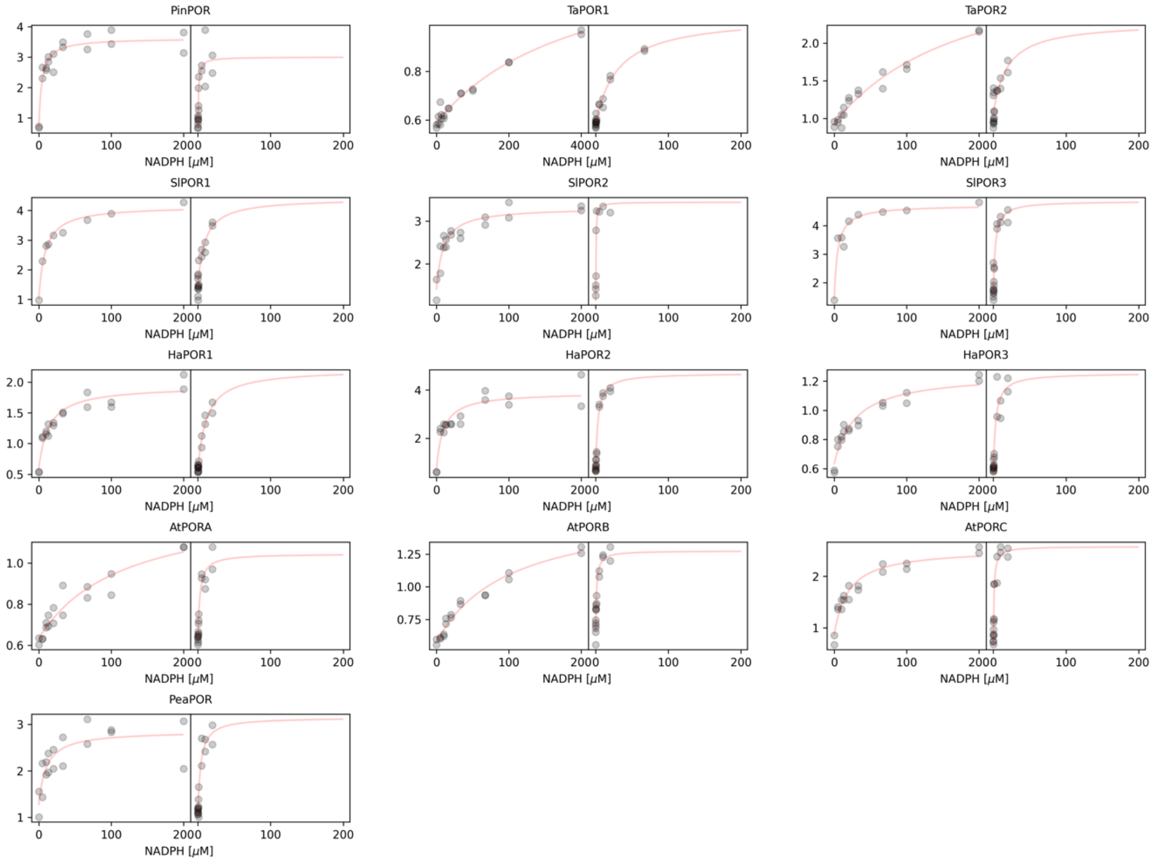


**Figure S3.** Relationship between the intensity ratio of 647/635 (left panels) and 655/635 (right panels) and NADPH concentration for different isoforms. The ratios are calculated to the spectra presented in Fig. S2. A fit of a modified Michaelis-Menten equation is shown.

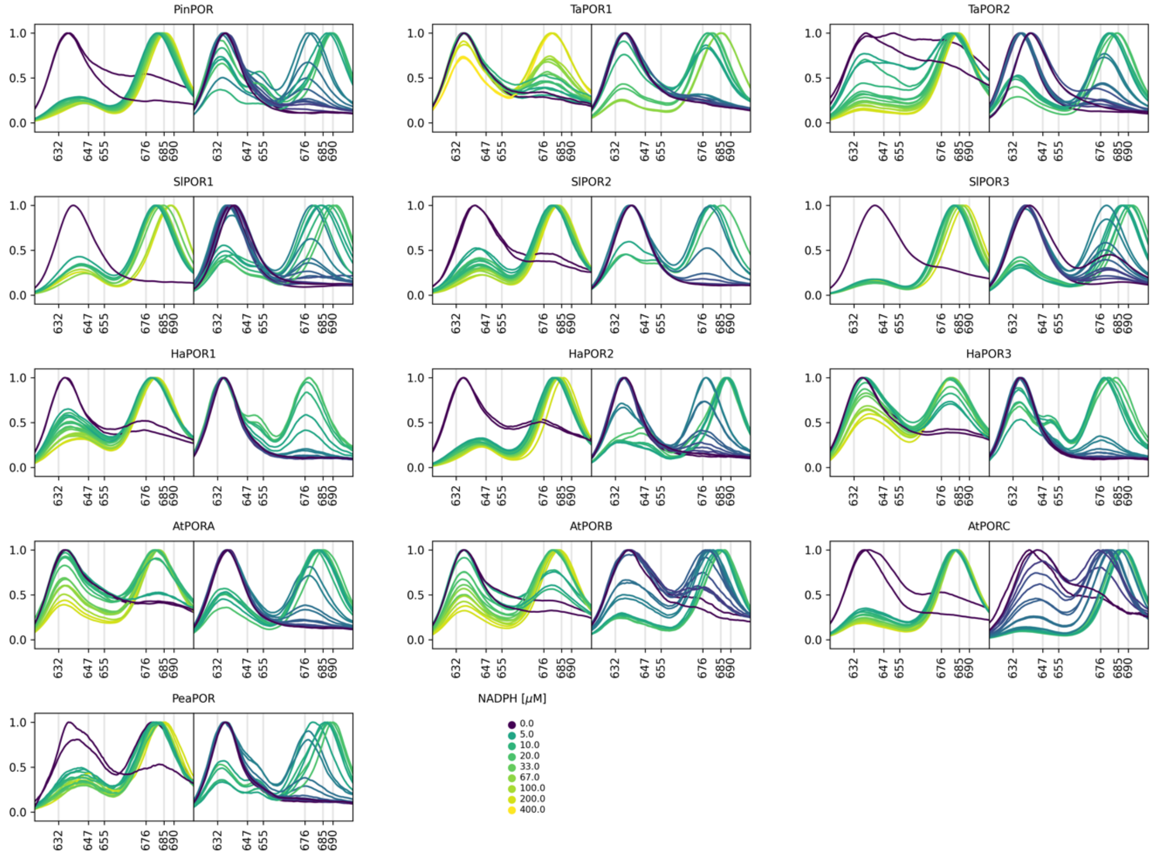


**Figure S4.** The spectra after 20 seconds of illumination of reaction mixtures for $K_M^L$ determination without (left panels) and with 400 µM OPT (right panels) for different isoforms. The reaction mixtures contained 15 µM LPOR, 5 µM pigment and variable NADPH concentrations.

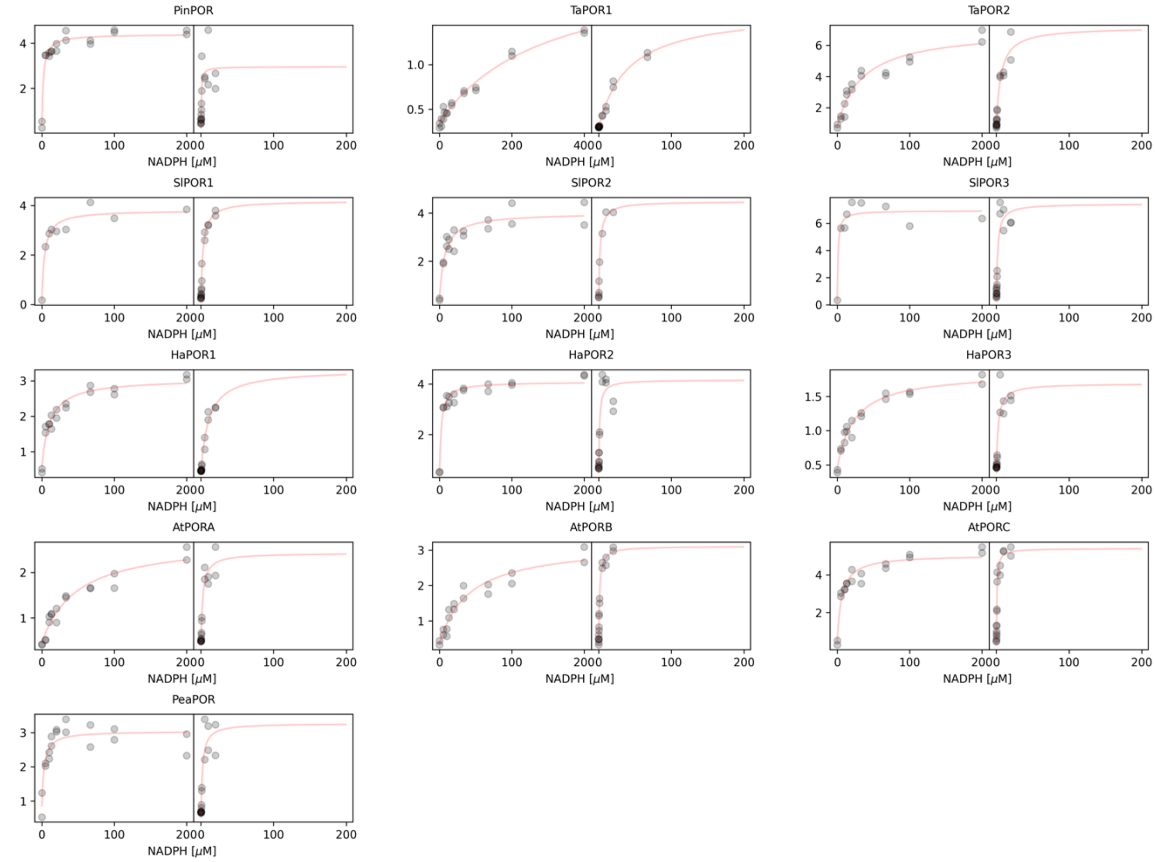


**Figure S5.** Relationship between Chlide/Pchlide and NADPH concentration for samples without (left panels) and with 400 µM OPT (right panels) for different isoforms for different isoforms. The ratios are calculated to the spectra presented in Fig. S4. A fit of a modified Michaelis-Menten equation is shown.

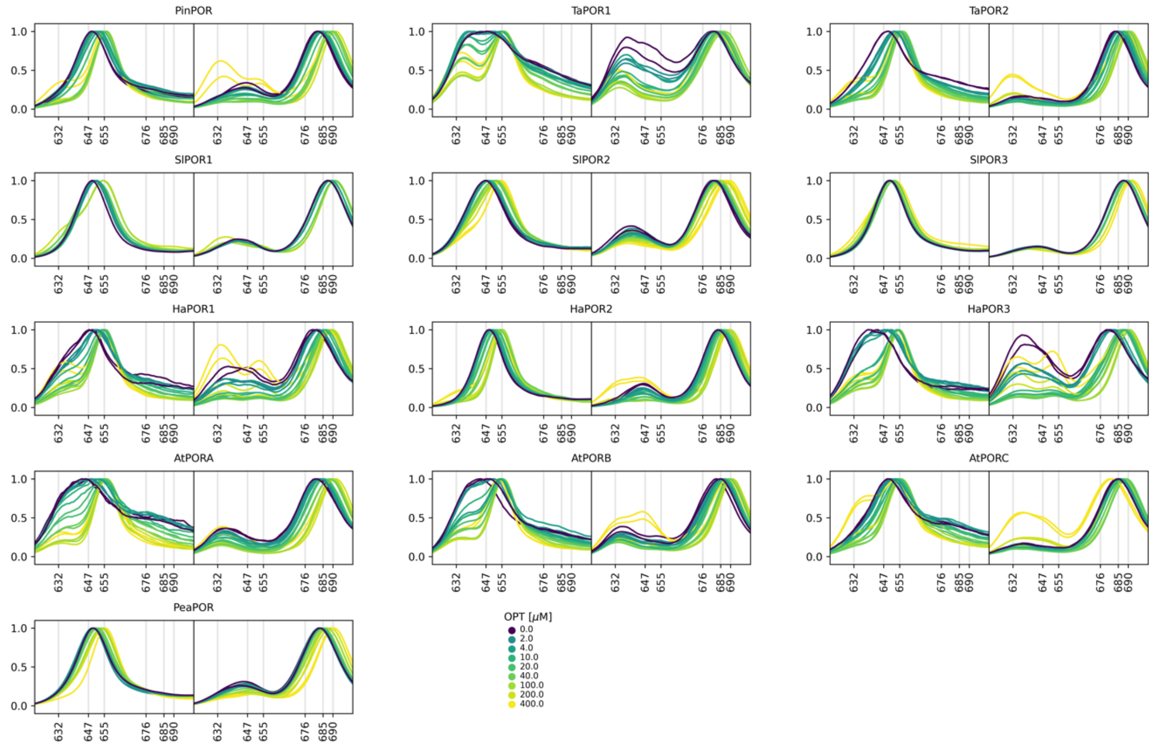


**Figure S6.** The spectra before (left panels) and after (right panels) 20 seconds of illumination of reaction mixtures with variable OPT concentrations for different isoforms. The reaction mixtures contained 15 μM LPOR, 5 μM pigment and 200 μM NADPH concentrations.

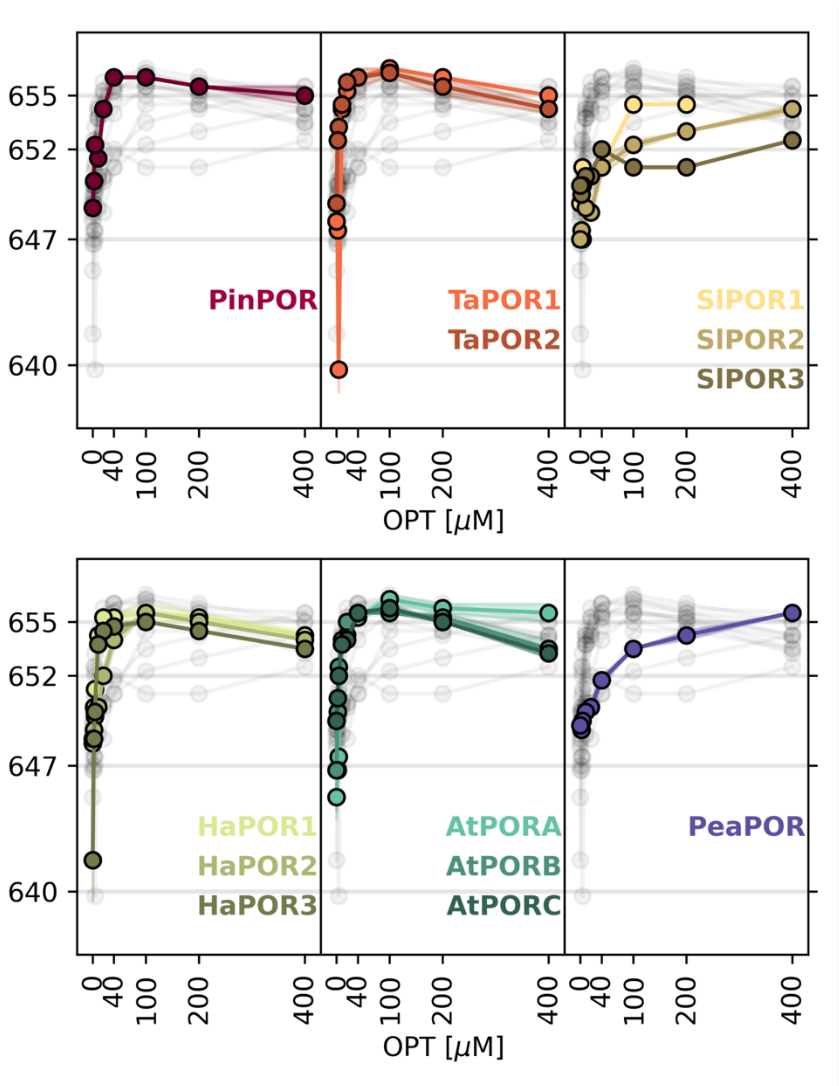


**Figure S7.** The relation between OPT concentration and emission maximum of Pchlide before the illumination for different isoforms. The maxima were read from the spectra in Fig. S6.

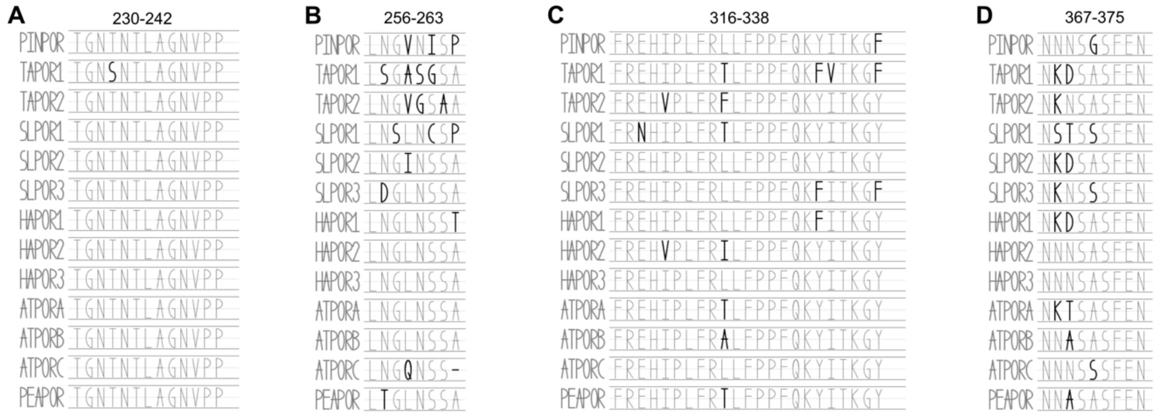


**Figure S8.** The sequences of selected region of LPOR. A. Pchlide loop. B. Oligomerization interface I. C. Helix α10. D. Oligomerization interface II.

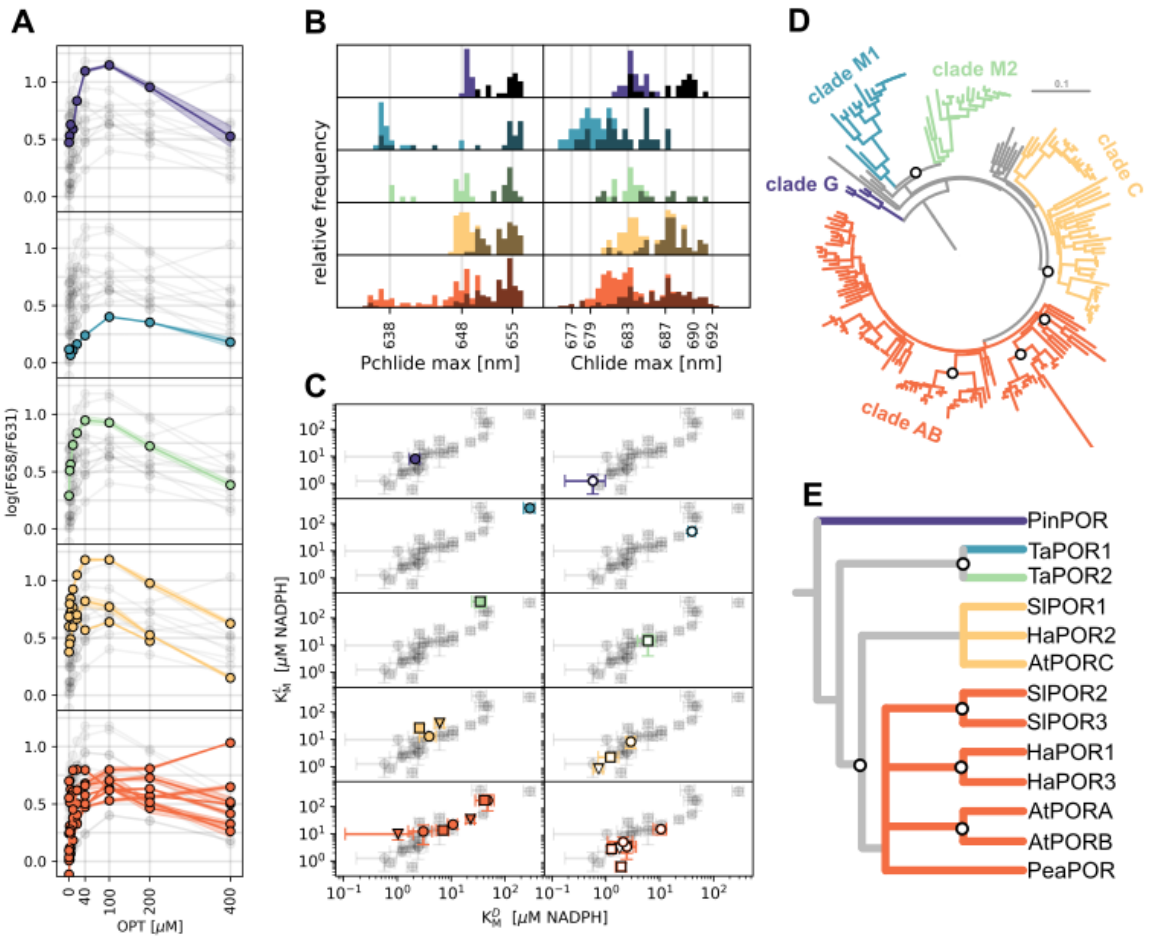


**Figure S9. Comparison of LPOR properties across clades.** (A) Relationship between the base-10 logarithm of the intensity ratio (658/631, oligomerization index) and OPT concentration. (B) Distribution of emission maxima for Pchlide and the corresponding Chlide maxima generated during the reaction. (C) Relationship between $K_M^D$ and $K_M^L$ within each clade. (D) Rooted phylogenetic tree of LPOR with clades marked; gene duplications are indicated by white circles. (E) Simplified phylogenetic tree showing the distribution of the studied isoforms within each clade. The data presented here corresponds to those in Fig. 2C, Fig. 4C, and Fig. 5B, but is rearranged according to clade affiliation. Shaded areas represent the standard deviation among replicates, and the overall data distribution is depicted in gray.

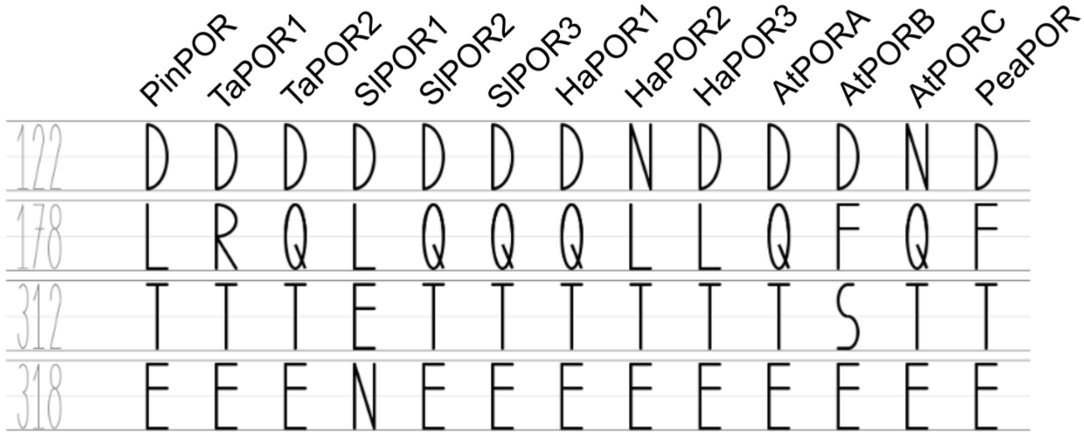


**Figure S10.** The residues at selected positions (122,178, 312 and 318) in different isoforms.

**Table S1**. The determined values of $K_M^D$ and $K_M^L$ constants presented in Fig. 3D.

| | dark | | | | illuminated | | | |
|---|---|---|---|---|---|---|---|---|
| | no lipids | | 400 uM OPT | | no lipids | | 400 uM OPT | |
| | $K_M^D$ | error | $K_M^D$ | error | $K_M^L$ | error | $K_M^L$ | error |
| PinPOR | 4.3 | 1.1 | 1.3 | 0.8 | 2.2 | 0.5 | 0.6 | 0.4 |
| TaPOR1 | 394.4 | 116.7 | 45.8 | 6.3 | 291.8 | 68.9 | 39.7 | 5.8 |
| TaPOR2 | 197.1 | 72.0 | 10.6 | 4.9 | 34.9 | 10.4 | 6.1 | 2.2 |
| SlPOR1 | 8.6 | 1.7 | 7.2 | 3.5 | 3.9 | 1.3 | 2.9 | 0.5 |
| SlPOR2 | 9.3 | 2.8 | 0.7 | 0.2 | 7.2 | 1.9 | 1.9 | 0.4 |
| SlPOR3 | 4.9 | 1.8 | 3.0 | 0.8 | 1.0 | 0.9 | 1.8 | 0.7 |
| HaPOR1 | 14.1 | 3.7 | 16.9 | 5.6 | 10.8 | 2.2 | 10.5 | 2.9 |
| HaPOR2 | 9.4 | 3.1 | 2.2 | 0.7 | 2.6 | 0.5 | 1.2 | 0.5 |
| HaPOR3 | 27.2 | 7.2 | 3.9 | 2.2 | 23.1 | 4.2 | 2.4 | 1.2 |
| AtPORA | 116.0 | 57.2 | 3.2 | 1.3 | 47.9 | 14.7 | 2.5 | 0.9 |
| AtPORB | 103.2 | 24.2 | 2.3 | 0.5 | 41.7 | 13.4 | 1.3 | 0.1 |
| AtPORC | 20.1 | 4.9 | 0.8 | 0.2 | 6.2 | 1.3 | 0.7 | 0.2 |
| PeaPOR | 10.8 | 6.1 | 4.7 | 2.1 | 3.0 | 1.4 | 2.1 | 1.0 |

**Table S2.** Primers used for gene cloning.

| gene | primer | primer sequence | Tm [C] |
|---|---|---|---|
| SlPOR1 | F | CCGCGCGGCAGCCATACAGCTGCTACCACTCCTGC | 70 |
| | R | TGTTAGCAGCCGGATTAAGCCAGTCCAACGAGCTTCTC | |
| SlPOR2 | F | CCGCGCGGCAGCCATACAATGGTTGCTTCTCCTGG | 66 |
| | R | TGTTAGCAGCCGGATTAAGCCAAACCAACGAGTTTCTCAC | |
| SlPOR3 | F | CCGCGCGGCAGCCATACAATGGTTGCATCTCCCG | 64 |
| | R | TGTTAGCAGCCGGACTAAGCCAATCCCACGAG | |
| PinPOR | F | CCGCGCGGCAGCCATACTGTGGCCGCACCAGTG | 71 |
| | R | TTGTTAGCAGCCGGATCAAGCAAGTCCAACAAGCTTTTCGC | |
| PeaPOR | F | TGCCGCGCGGCAGCCATACAGCGGCTCCGGCCACTC | 71 |
| | R | GCTTTGTTAGCAGCCGGATTAGGCCAAACCAACAAGCTTCTC | |
| HaPOR1 | F | GTGCCGCGCGGCAGCCATGTAGCCACTACAAGTACTCCTCC | 69 |
| | R | GGGCTTTGTTAGCAGCCGGATCAAGCCAACCCAACAATCTTC | |
| HaPOR2 | F | GTGCCGCGCGGCAGCCATCAAACAGCAACCGTAGCTC | 67 |
| | R | GGGCTTTGTTAGCAGCCGGACTAAGCCAAACCAACAAGCTTC | |
| HaPOR3 | F | GTGCCGCGCGGCAGCCATGCTGTAGCCACAACTCCAG | 69 |
| | R | GGGCTTTGTTAGCAGCCGGATTAAGCCAACCCGACCAAC | |
| pET15b | F | TCCGGCTGCTAACAAAGCCCGA | 71 |
| | R | ATGGCTGCCGCGCGGCAC | |

**Table S3.** Range definition for Pchlide loop, oligomerization interface I, helix α10, and oligomerization interface II.

| enzyme | residues range | | sequence |
|---|---|---|---|
| PinPOR | 228 | 240 | TGNTNTLAGNVPP |
| | 254 | 261 | LNGVNISP |
| | 314 | 336 | FREHIPLFRLLFPPFQKYITKGF |
| | 365 | 373 | NNNSGSFEN |
| TaPOR1 | 216 | 228 | TGNSNTLAGNVPP |
| | 242 | 249 | LSGASGSA |
| | 303 | 325 | FREHIPLFRTLFPPFQKFVTKGF |
| | 354 | 362 | NKDSASFEN |
| TaPOR2 | 246 | 258 | TGNTNTLAGNVPP |
| | 272 | 279 | LNGVGSAA |
| | 332 | 354 | FREHVPLFRFLFPPFQKYITKGY |
| | 383 | 391 | NKNSASFEN |
| SlPOR1 | 228 | 240 | TGNTNTLAGNVPP |
| | 254 | 261 | LNSLNCSP |
| | 314 | 336 | FRNHIPLFRTLFPPFQKYITKGY |
| | 365 | 373 | NSTSSSFEN |
| SlPOR2 | 224 | 236 | TGNTNTLAGNVPP |

| | | | |
|---|---|---|---|
| | 250 | 257 | LNGINSSA |
| | 310 | 332 | FREHIPLFRLLFPPFQKYITKGY |
| | 361 | 369 | NKDSASFEN |
| SlPOR3 | 226 | 238 | TGNTNTLAGNVPP |
| | 252 | 259 | LDGLNSSA |
| | 312 | 334 | FREHIPLFRLLFPPFQKFITKGF |
| | 363 | 371 | NKNSSSFEN |
| HaPOR1 | 227 | 239 | TGNTNTLAGNVPP |
| | 253 | 260 | LNGLNSST |
| | 313 | 335 | FREHIPLFRLLFPPFQKFITKGY |
| | 364 | 372 | NKDSASFEN |
| HaPOR2 | 216 | 228 | TGNTNTLAGNVPP |
| | 242 | 249 | LNGLNSSA |
| | 302 | 324 | FREHVPLFRILFPPFQKYITKGY |
| | 353 | 361 | NNNSASFEN |
| HaPOR3 | 213 | 225 | TGNTNTLAGNVPP |
| | 239 | 246 | LNGLNSSA |
| | 299 | 321 | FREHIPLFRLLFPPFQKYITKGY |
| | 350 | 358 | NNNSASFEN |
| AtPORA | 234 | 246 | TGNTNTLAGNVPP |
| | 260 | 267 | LNGLNSSA |
| | 320 | 342 | FREHIPLFRTLFPPFQKYITKGY |
| | 371 | 379 | NKTSASFEN |
| AtPORB | 230 | 242 | TGNTNTLAGNVPP |
| | 256 | 263 | LNGLNSSA |
| | 316 | 338 | FREHIPLFRALFPPFQKYITKGY |
| | 367 | 375 | NNASASFEN |
| AtPORC | 231 | 243 | TGNTNTLAGNVPP |
| | 257 | 263 | LNGQNSS |
| | 316 | 338 | FREHIPLFRLLFPPFQKYITKGY |
| | 367 | 375 | NNNSSSFEN |
| PeaPOR | 228 | 240 | TGNTNTLAGNVPP |
| | 254 | 261 | LTGLNSSA |
| | 314 | 336 | FREHIPLFRTLFPPFQKYITKGY |
| | 365 | 373 | NNASASFEN |

**Video S1.** The animated morph transitions between the predicted AtPORB conformations and the reference structure 7JK9.
